# Large-scale duplication events underpin population-level flexibility in tRNA gene copy number in *Pseudomonas fluorescens* SBW25

**DOI:** 10.1101/2022.12.02.516541

**Authors:** Zahra Khomarbaghi, Wing Y. Ngan, Gökçe B. Ayan, Sungbin Lim, Gunda Dechow-Seligmann, Pabitra Nandy, Jenna Gallie

**Author notes:** Joint authors: these authors contributed equally to this work (Zahra Khomarbaghi, Wing Y. Ngan). Zahra Khomarbaghi, Division of Evolution and Genomic Sciences, School of Biological Sciences, Faculty of Biology, Medicine and Health, University of Manchester, Manchester, M13 9PL, United Kingdom.

## Abstract

The complement of tRNA genes within a genome is typically considered to be a (relatively) stable characteristic of an organism. Here we demonstrate that bacterial tRNA gene set composition can be more flexible than previously appreciated, particularly regarding tRNA gene copy number. We report the high-rate occurrence of spontaneous, large-scale, tandem duplication events in laboratory populations of the bacterium *Pseudomonas fluorescens* SBW25. The identified duplications are up to ∼1 Mb in size (∼15 % of the wildtype genome) and are predicted to change the copy number of up to 917 genes, including several tRNA genes. The observed duplications are inherently unstable: they occur, and are subsequently lost, at extremely high rates. We propose that this unusually plastic type of mutation provides a mechanism by which tRNA gene set diversity can be rapidly generated, while simultaneously preserving the underlying tRNA gene set in the absence of continued selection. That is, if a tRNA set variant provides no fitness advantage, then high-rate segregation of the duplication ensures the maintenance of the original tRNA gene set. However, if a tRNA gene set variant is beneficial, the underlying duplication fragment(s) may persist for longer and provide raw material for further, more stable, evolutionary change.

## INTRODUCTION

The complement of tRNA genes encoded by an organism (the ‘tRNA gene set’) is an important determinant of translational efficiency (1, 2) and disease trajectories (3–5). In bacteria, tRNA gene set composition varies considerably, yet the tRNA gene set of any given bacterial strain is typically treated as a stable entity. For example, the tRNA gene sets of *Pseudomonas fluorescens* strains SBW25 and Pf0-1 are reported to respectively contain 66 and 73 canonical tRNA genes, encoding 39 and 40 distinct species of mature tRNA (each with a unique anticodon-amino acid combination) (6–8). In this work we highlight that, at the population level, the tRNA gene set of a bacterial strain may be more flexible than previously appreciated, particularly with regard to the number of gene copies encoding each type of tRNA.

The composition of tRNA gene sets differs across the bacterial phylogeny; extant bacterial tRNA gene sets differ in both the tRNA species that they encode, and in the number of gene copies coding for each tRNA species (7, 9–15). Variations in tRNA species encoded are possible because of wobble base pairing, the process by which some tRNAs can – either naturally (16) or through post-transcriptional modifications (17, 18) – translate multiple codons. Ultimately, wobble base pairing allows all 61 sense codons to be translated by far fewer than the number of directly corresponding tRNA species, with most bacteria encoding between 25 and 46 different types of tRNA (7, 9, 19). In contrast to this apparently reductionist strategy, the tRNA species that are present within a bacterial genome may be encoded by multiple – and often, identical – gene copies (7, 10).

The content of a tRNA gene set has a strong influence on the composition of the mature tRNA pool, which in turn is a major determinant of the speed and accuracy of translation (1, 2, 20). The capacity of a tRNA set for efficient translation depends on a number of additional factors, many of which co-evolve with tRNA set composition. Prominent examples include (i) which post-transcriptional tRNA modification enzymes are present (affecting anticodon-codon matching patterns) (15, 18, 21, 22), (ii) codon usage (which influences the demand for specific tRNAs) (23–29), and (iii) environmental conditions that affect anticodon-codon base pairing properties (*e.g.*, pH, temperature) (30, 31). Evolutionary optimization of translational efficiency is expected to shape the evolution of each of these factors, particularly under conditions where efficient translation is most important (*e.g.*, during rapid bacterial growth) (32–35). For example, bacteria with faster growth rates tend to carry tRNA gene sets with fewer tRNA species, encoded by higher gene copy numbers (11). Empirically, multiple copies of several *Escherichia coli* tRNA genes have been demonstrated to be advantageous only under conditions that support rapid growth (and hence, rapid translation) (36).

The molecular mechanisms by which tRNA genes are gained and lost from tRNA gene sets have been inferred by phylogenetic analyses. Such computational studies have provided evidence for the evolution of extant bacterial tRNA gene sets via a number of routes, including: (i) mutation of tRNA genes to encode different tRNA species (‘anticodon switching’), (ii) loss of tRNA genes by deletion, and (iii) acquisition of new tRNA genes, either by within-genome duplication events, or by horizontal gene transfer (10, 12–14, 37–39). In addition, we recently provided an empirical example of a bacterial tRNA gene set undergoing adaptive evolution; elimination of tRNA-Ser(CGA) from the mature tRNA pool of the fast-growing, model bacterium *P. fluorescens* SBW25 was compensated by within-genome duplication events that serve to increase the copy number of the tRNA gene *serTGA* (and corresponding levels of tRNA-Ser(UGA) in the mature tRNA pool) (6). Others have reported that translational challenges imposed by (i) novel codon usage patterns, or (ii) the fitness costs associated with antibiotic resistance mutations can be readily alleviated by similar duplication (and, sometimes, downstream amplification) events that affect the copy numbers of remaining tRNA genes (40, 41). Together, these studies demonstrate a considerable degree of evolutionary flexibility in the composition of bacterial tRNA gene sets.

In the above-mentioned study of the tRNA gene set of *P. fluorescens* SBW25 (6), the adaptive increase in *serTGA* copy number was shown to be mediated through the tandem duplication of large sections of the *P. fluorescens* SBW25 chromosome (ranging in size from ∼45 kb to ∼290 kb). In addition to the 91 bp target gene (*serTGA*), these duplication events affect the dosage of up to a further 278 genes, and generate considerable alterations in chromosome size and architecture. Similar types of large-scale, tandem duplications are a pervasive phenomenon in bacterial genomes (42–46) and – despite their large size and widespread gene dosage effects – they can provide a net selective advantage in response to a range of selective pressures (47–53). Notably, these types of mutations have been reported to arise spontaneously, and at extremely high rates; estimates range from frequencies of 10^-1^ to 10^-5^ duplications per cell per generation (43, 54–57), several orders of magnitude higher than those expected for single nucleotide polymorphisms (SNPs) (58). Given their pervasiveness and ease of occurrence, such large-scale, tandem duplications challenge the prevailing view of fixed, stable tRNA gene sets, particularly in large, rapidly growing bacterial populations.

In this work, we firstly show that large-scale, tandem duplication events affecting the *P. fluorescens* SBW25 tRNA gene content are not limited to the genomic region surrounding *serTGA* (located at ∼4.16 Mb on the SBW25 chromosome). Specifically, we demonstrate that a fitness defect caused by deleting a locus containing four tRNA genes (two copies of each of *gluTTC* and *glyGCC*) is repeatedly and rapidly compensated by large-scale, tandem duplication events encompassing the remaining *glyGCC* gene (at genomic position ∼2.38 Mb). Secondly, we provide empirical evidence that these duplication fragments arise, and are lost, at very high frequencies; we show that in each independently evolving mutant population, multiple distinct duplication fragments (and hence, tRNA gene sets) rapidly arise and compete.

## MATERIALS AND METHODS

### Strains and growth conditions

A list of strains is provided in Supplementary Table S1; construction details can be found in Supplementary Text S1. Unless otherwise stated, *P. fluorescens* SBW25 was grown in King’s Medium B (KB) (59) for ∼16 hours at 28°C with shaking. For growth curves, strains were grown in either liquid KB, or minimal (M9) medium (60) supplemented with 4 % (w/v) glucose. During engineering procedures, *E. coli* strains were grown in Lysogeny Broth (LB) for 16-18 hours at 37°C with shaking.

### Photography

Colonies were grown for the time indicated, at room temperature (∼20-22°C) or 28°C, on KB agar. Colonies were visualized under a Leica MS5 dissection microscope and photographed with a VisiCam® 1.3 (VWR International). Photographs were cropped and, where noted in figure legends, the exposure and/or brightness uniformly altered in Preview (v11.0) to improve visibility.

### Genotype growth curves

Seven colonies per genotype were inoculated into individual wells of a 96-well pre-plate (each containing 200 μL KB), and grown overnight (28°C, 200 rpm). From each well, 2 μL of grown culture was transferred into 198 μL fresh KB, covered with a Breathe-Easy® membrane, and grown at 28°C in an Epoch 2 Microplate Spectrophotometer (Agilent BioTek). Absorbance at 600nm (OD_600_) was measured at five-minute intervals, with 5 seconds of 3 mm orbital shaking preceding each read. Gen5^TM^ Data Analysis Software (version 3.00.19) was used to calculate the maximum growth rate (the calculated value of the mean slope, V_max_) and lag time (time interval between the line of maximum slope of the propagation phase and the absorbance baseline at time zero) for each growth curve, with the following parameters: calculation window=1-8 hours (KB) or 1-10 hours (M9); sliding time interval per calculation (“*n*”)=6 points). The data was then exported to Excel for further processing and drawing of graphs (Figures 3 and 4). Where assumptions are satisfied, parametric one-way ANOVA was used to test for differences in the mean maximum growth rate between pairs of genotypes. Where assumptions are violated, non-parametric Dunn’s tests, followed by the Benjamini-Hochberg procedure to correct for multiple comparisons, were used to test for differences in the median lag time between pairs of genotypes.

### Competition assays

One-to-one competition experiments were used to determine the relative fitness of 9 genotype pairs under conditions similar to those of a single growth cycle of the evolution experiment (see below). Competitions were performed in two blocks, with between 5 and 8 replicates per pair of genotypes. Single colonies of each genotype were grown on KB agar (48 hours, room temperature) and, for each competition replicate, a single colony of each competitor was inoculated into an overnight culture (4 mL KB, 28°C, 200 rpm). Once grown, the two competitors for each competition were mixed in a ∼1:1 ratio. Ten μL of each mixture was used to inoculate 10 mL KB in a 50 ml Falcon tube, and grown for 24 hours (28°C, 200 rpm). Cell samples taken from each competition tube at the start (T_0_) and end (T_24_) of the incubation period were serially diluted and plated on LB agar containing 60 μg mL^-1^ X-gal (48 hours, room temperature). The number of colonies per competitor were recorded at T_0_ and T_24_, where differentiation between competing genotypes was achieved through distinct colony morphologies and/or colours (in competitions including neutrally marked SBW25*-lacZ* (61)). The change in the ratio of the competitors was calculated, and relative fitness of the competitors determined (62). Parametric one-sample *t*-tests or, where normality assumptions were violated, non-parametric Wilcoxon rank sum tests, were used to detect statistical differences in the mean relative fitness of genotype pairs. The Bonferroni procedure was subsequently used to correct for multiple comparisons.

### Evolution experiment

SBW25 (wildtype) and ΔEGEG (quadruple tRNA gene deletion mutant) were streaked from glycerol stocks onto KB agar and grown (48 hours, 28°C). Five single colonies per genotype were grown overnight (4 mL KB; 28°C, 200 rpm). Grown cultures (“day 0”) were vortexed and 100 μL of each used to inoculate 9.9 mL KB in a 50 mL Falcon tube (28°C, shaking, 24 hours). Every 24 hours thereafter, 1 % (100 μL) of each culture was transferred to a fresh 9.9 mL KB, and a sample of the population frozen in glycerol saline at −80°C. The experiment was continued until day 28. The later days (∼day 21 onwards) of mutant lines M4 and M5 were discovered to contain external contaminants (see Supplementary Figure S1), and so were removed from downstream population analyses (*i.e.*, Figure 7 and Figure 8). Extensive checks for external and cross contamination events were performed on the remaining 8 lines (including day 28 population-level whole genome re-sequencing, population colony morphology checks, and population growth curve analyses).

### Population growth curves

Forty populations of interest (10 evolving lines, at each of 4 time points) were retrieved from glycerol stocks by re-suspending large scrapings in 200 μL sterile Ringer’s solution. Each re-suspension was used to inoculate 4 wells of 96-well pre-plates, and grown overnight (*i.e.*, 4 technical replicates; 2 μL re-suspension into 198 μL KB per replicate). Subsequently, grown pre-cultures were transferred to a new 96-well plate (2 μL pre-culture into 198 μL fresh KB), and the protocol continued as described above for the genotype growth curves. This protocol was repeated three times per population (*i.e.*, 3 biological replicates), giving a total of 12 replicates per population of interest (4 technical replicates x 3 biological replicates). The final population growth curves are presented in Supplementary Figure S1.

### Genotype whole genome re-sequencing

Six single genotypes (W1-1, M1-1, M2-1, M3-1, M4-1, M5-1) were purified from day 21 of the evolution experiment, including 1 genotype from each of the 5 mutant lines (M1-M5), and 1 from a representative wildtype control line (W1). Genomic DNA was isolated from 1 mL of overnight culture of each genotype (DNeasy Blood & Tissue Kit; Qiagen). Paired-end, 250-bp reads were generated with an Illumina MiSeq instrument at The Max Planck Institute for Evolutionary Biology (Ploen, Germany), via standard procedures. Raw reads are available at NCBI Sequence Read Archive (SRA accession number: PRJNA790786) (63). A minimum of 1.08 million raw reads per genotype were obtained and aligned to the SBW25 reference genome sequence (NC_012660.1) (8) using *breseq* (64–66) and Geneious Prime (v2023.2.1). A minimum mean coverage of 36.4 reads per bp was obtained. Per genotype, a list of the raw differences predicted by *breseq* was manually curated to give a list of final mutation predictions (see Supplementary Table S2 for lists and further details). Each mutation prediction was confirmed by standard PCR or duplication junction PCR (see below) and, where indicated, Sanger sequencing (see Supplementary Text S2).

### Mature tRNA pool sequencing (YAMAT-seq)

Three independent replicates of 8 genotypes (*i.e.*, 24 samples) were used for YAMAT-seq (67) according to the method described in (6). Briefly, (i) per sample, a single colony was chosen from KB agar (in the case of genotypes carrying duplication fragments, rapidly-growing colonies were chosen in an attempt to mitigate the inherent instability of the duplication fragment; see stability test method below), grown in overnight KB culture, diluted 1/10 in fresh KB, and grown to mid-exponential phase, (ii) total RNA was isolated from each rapidly-growing liquid culture using a TRIzol Max Bacterial RNA isolation kit, (iii) amino acids were removed from mature tRNAs by deacylation treatment with 20 mM Tris-HCl, (iv) Y-shaped, DNA/RNA hybrid adapters (Eurofins; see Supplementary Table S1) were annealed to the conserved, exposed 5’-NCCA-3’ and 3’-inorganic phosphate-5’ ends of non-acylated tRNAs, and ligated T4 RNA ligase 2, (v) ligation products were reverse-transcribed into cDNA using SuperScript^TM^ III reverse transcriptase, (vi) cDNA products were amplified, and a unique barcode added, with eleven cycles of PCR using Phusion^TM^, (vii) products were quantified using a Bioanalyzer and fluorescent Nanodrop, and mixed in equimolar amounts, (viii) the mixed sample was run on a 5 % polyacrylamide gel, the bands between ∼210 bp and ∼280 bp (tRNAs plus adapters and barcodes) cut out and co-extracted overnight (by soaking in deionized, ultrapure water (Illumina)), (ix) the extracted liquid sample was collected by centrifugation through a filtered microcentrifuge tube (cellulose acetate membrane with pore size 0.45 μm; Merck CLS-8162), (x) the purified product was quantified and deep sequenced at the Max Planck Institute for Evolutionary Biology (Ploen, Germany). Single-end, 150 bp reads were generated with a NextSeq 550 Output v2.5 kit (Illumina) (NCBI Gene Expression Omnibus accession number GSE196705) (68).

### YAMAT-seq analysis

The raw YAMAT-seq sequencing reads were split into 24 samples by extracting exact matches to each unique, 6-bp Illumina index. The resulting 24 set of raw reads, each containing a minimum of 1,470,795 reads, were analysed using Geneious Prime (version 11.1.4). Reads of the expected length (80-151 bp) were extracted (resulting in at least 99.8 % read retention per sample). The extracted reads were aligned to a set of 42 reference tRNA sequences from SBW25 (see Supplementary Text S4) (Geneious settings: per read, up to 10 % mismatches, gaps of <3 bp, up to five ambiguities; discard reads that align equally well to more multiple reference sequences). The unused reads for each sample were *de novo* assembled, and the resulting contigs were checked to ensure that none contained substantial numbers of tRNA reads. The within-sample proportion of reads aligned to each tRNA species was calculated. Next, mean mature tRNA proportions were calculated for each genotype across the 3 replicates (Supplementary Table S4). DESeq2 (69) was used in R (version 3.6.0) to detect tRNA expression differences between pairs of genotypes (Supplementary Table S5). DESeq2 corrects for multiple testing (Benjamini-Hochberg procedure) (70).

### Stability test

Duplication fragment stability was estimated using colony morphology stability tests, according to the method detailed in (71). Briefly, 10 genotypes (WT, rWT, ΔEGEG, Δ*glyGCC*, W1-1, M1-1, M2-1, M3-1, M4-1, and M5-1) were streaked from glycerol stocks onto KB agar plates and grown (48 hours, 28°C). Five colonies per genotype were used to inoculate 4 mL KB cultures (28°C, 200 rpm, overnight). For duplication-carrying genotypes, all chosen colonies were large (*i.e.*, were expected to retain the duplication). Cultures were then dilution plated on KB agar (48 hours, room temperature), and the number of large and small colonies per plate was enumerated. Between 41 and 295 colonies were recorded per replicate, with no more than ∼200 colonies counted on any given plate. For each replicate, the proportion of small colonies was calculated. A non-parametric Kruskal-Wallis test was used to test for evidence of a difference in the median of at least 1 founder (*p*=6.25e-07). To determine which genotypes showed a different median rate of small colony enumeration as compared with SBW25 (wildtype), Dunn’s test was used with Benjamini-Hochberg procedure to adjust for multiple comparisons. A Welch non-parametric two-sample *t*-test was used to test for a difference in the mean proportion in small colonies arising from the genotype carrying the small duplication fragment (M3-1), and those carrying a large duplication fragment (M1-1, M2-1, M4-1, M5-1).

### Duplication junction PCRs

Duplication junction PCRs were used in this work to (i) confirm the computationally-predicted duplications in genotypes isolated from day-21 of the evolution experiment (see Supplementary Text S2), and (ii) reveal the evolutionary trajectories of duplication fragments in evolving populations (method described here). For each trajectory of interest, 3 independent rounds of duplication junction PCR were performed. Per round, PCR templates were prepared as follows: a large chunk of the relevant glycerol stock was melted at room temperature, and 10 μL used to inoculate a 4 mL KB culture (20 h at 28°C, 200 rpm). Genomic DNA was extracted from 1 mL of culture (DNeasy Blood & Tissue Kit; Qiagen) and quantified on a Nanodrop. Each genomic DNA sample was used as a template for the following PCRs, at a final concentration of 1 ng μL^-1^: (PCRs 1 and 2) the duplication junction PCR(s) of interest, targeting ∼1 Mb duplications (with primer pair M1and2junct_f/M5_junct_r), and/or ∼0.49 Mb duplications (with primer pair M3_junct_f/r); (PCR 3) a standard, control PCR targeting a genomic region unaffected by the duplication events (at genomic position ∼0.01 Mb; primer pair *glyQ_*f_/r). PCRs were performed with GoTaq Flexi (Promega) for 30 cycles of: 30 sec at 95°C, 30 sec at 58°C or 59°C, 72°C for 1 min per kb of expected product. Five μL of each PCR product was mixed with loading dye and run on a 1 % agarose gel (SYBR Safe; Life Technologies) at 90 V for ∼45 min, against 1 kb DNA ladder (Promega) and photographed under UV illumination (see Supplementary Figures S3 and S4 for examples). PCR product intensities were quantified using ImageJ2 (version 2.14.0/1.54f) (72), and duplication junction PCR intensities were normalized using the corresponding control PCR (see Supplementary Table S6). Final evolutionary trajectories are plotted in Figure 7 and Supplementary Figure S5A.

### Population whole genome re-sequencing

Nine samples were used for population-level whole genome re-sequencing: 8 day-28 populations (from lines M1-M3 and W1-W5), and the non-evolved SBW25 wildtype genotype (which, as a control, was treated as a population sample in downstream computational analyses). For each of the 8 day-28 samples, the corresponding day-27 glycerol stock was grown for 1 cycle of the evolution experiment (in 9.9 mL KB: 180 μL thawed glycerol stock, consisting of ∼100 μL cell culture and ∼80 μL glycerol; 28°C, 200 rpm, 24 hours). For the control sample, a single SBW25 colony picked from KB agar was grown under the same conditions. Genomic DNA was extracted from 2 mL of each of the nine overnight cultures (DNeasy Blood & Tissue Kit; Qiagen). Library preparation and sequencing was performed by Novogene Europe (Cambridge, UK), using Illumina technology on a NovaSeq 6000 platform. The 150-bp, paired-end raw reads are available at NCBI Sequence Read Archive (SRA accession number: PRJNA1035442) (63). Raw reads were quality-trimmed using fastp (settings: --detect_adapter_for_pe, --cut_right, minimum read length 50 bp) (73). The polymorphism setting in *breseq* (64–66) was used to align each set of trimmed reads to the reference sequence (SBW25; NC_012660.1). This gave a minimum mean coverage of 154.9 reads per bp, and a set of predicted mutations in each sample (Supplementary Tables S2 and S7). Geneious Prime (v2023.2.1) was used to (i) confirm and extend the set of the duplication fragments predicted by *breseq* (Supplementary Text S3), and (ii) rigorously check for contamination in evolving populations (none was detected).

## RESULTS

### Severe reduction in essential tRNA species’ gene copy number slows growth

The *P. fluorescens* SBW25 genome is predicted to contain 66 canonical tRNA genes encoding 39 tRNA species, 14 of which are coded for by multiple (between two and five) gene copies (Figure 1A) (6, 8, 74). This work initially focusses on two such multicopy tRNA species: tRNA-Gly(GCC) (encoded by three identical *glyGCC* gene copies) and tRNA-Glu(UUC) (encoded by four identical *gluTTC* gene copies). Two copies of each gene are found together at chromosomal position ∼2.03 Mb (the *gluTTC-glyGCC-gluTTC-glyGCC* locus, hereafter referred to as “*EGEG*”) (Figure 1A-B). The third *glyGCC* copy is a lone tRNA gene at genomic position ∼2.38 Mb, while the final two *gluTTC* genes occur, together with two copies of *alaGGC*, at ∼4.65 Mb. Both of the focal tRNA species are essential: at least one functional copy of each of *glyGCC* and *gluTTC* is expected to be required for translation and growth. Specifically, tRNA-Gly(GCC) is predicted to be the sole decoder of codons 5’-GGC-3’ (cognate match) and 5’-GGU-3’ (G_34_:U_3_ wobble match), while tRNA-Glu(UUC) is required to translate 5’-GAA-3’ (cognate) and 5’-GAG-3’ (G_34_:U_3_ wobble) (Figure 1C).

**Figure 1.**
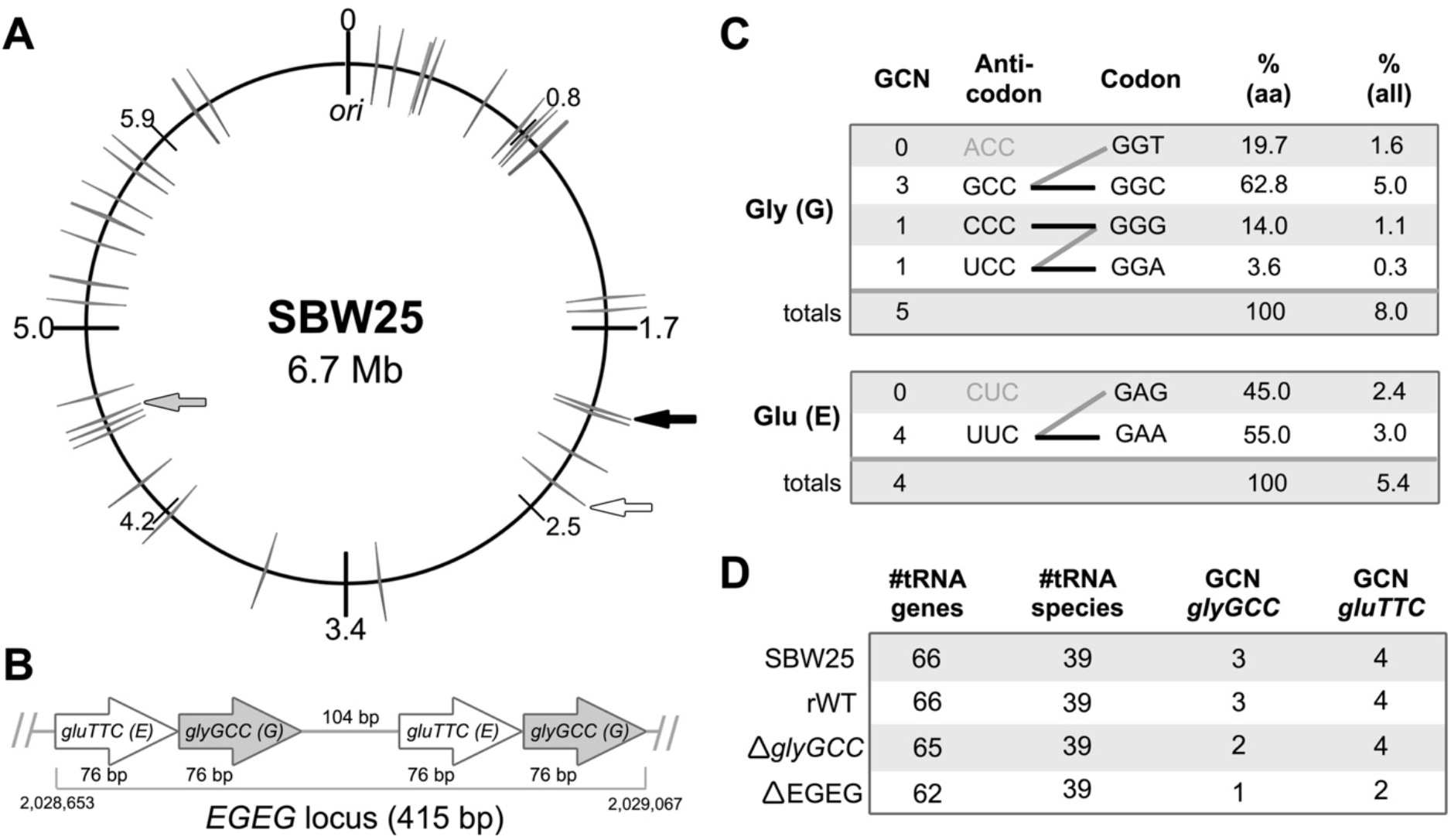
Translation of glycine (G) and glutamic acid (E) in *P. fluorescens* SBW25. (**A**) Cartoon of the SBW25 chromosome depicting loci predicted by GtRNAdb2.0 to encode canonical tRNAs. Grey lines=tRNA loci (each representing between one and four tRNA genes); black arrow=*gluTTC-glyGCC-gluTTC-glyGCC* (“*EGEG*”) tRNA locus deleted by genetic engineering; white arrow=*glyGCC* lone tRNA locus (encoding the remaining copy of *glyGCC*); grey arrow=*gluTTC-alaGGC-gluTTC-alaGGC* tRNA locus (encoding the remaining two copies of *gluTTC*). (**B**) Details of the *EGEG* tRNA locus indicated by the black arrow in panel A. (**C**) Putative translational relationships between anticodons and codons for glycine (top) and glutamic acid (bottom). Codons are listed with their cognate anticodons (black=tRNA species encoded in genome; grey=not encoded). Predicted anticodon-codon matching patterns are indicated by joining lines (black=cognate match; grey=wobble match). Percentage contribution of each codon to (i) the relevant amino acid (% aa), and (ii) all amino acids, is provided to 1 d.p. from GtRNAdb2.0 (tRNAscan-SE version 2.0.2; February 2019) (7). (**D**) Details of the tRNA gene sets carried by engineered genotypes.

We began by constructing two tRNA gene deletion genotypes in the SBW25 background: (i) ΔEGEG, in which the entire *EGEG* locus was removed from SBW25 (leaving two copies of *gluTTC* and only a single copy of *glyGCC*), and (ii) Δ*glyGCC*, in which the lone *glyGCC* copy at position ∼2.03 Mb was removed (leaving two copies of *glyGCC*, and all four copies of *gluTTC*) (Figure 1D; Supplementary Text S1). The two *gluTTC* copies at position ∼4.65 Mb were not initially targeted because the tRNA genes with which they co-localize are expected to be essential for survival (*i.e.*, they are the only two copies of *alaGGC*, encoding the essential tRNA-Ala(GGC)) (Figure 1A).

Deletion of *EGEG* results in an immediately obvious growth defect; compared with SBW25, ΔEGEG forms smaller colonies on KB and M9 agar plates, and grows more slowly in liquid culture (Figure 2). Further, wildtype-like growth is restored by the re-introduction of *EGEG* into ΔEGEG (generating the reconstructed wildtype, rWT). The observed reduction in growth is presumably due to a drop in levels of either tRNA-Glu(UUC), tRNA-Gly(GCC), or both. However, we note that there is at least partial redundancy among the *glyGCC* gene copies; removal of the lone *glyGCC* gene has no discernible effect on SBW25 growth under the conditions tested (Δ*glyGCC* in Figure 2). This observation is consistent with earlier reports that multiple copies of various tRNA genes (including *glyGCC*) show a degree of redundancy in *E. coli* (36).

**Figure 2.**
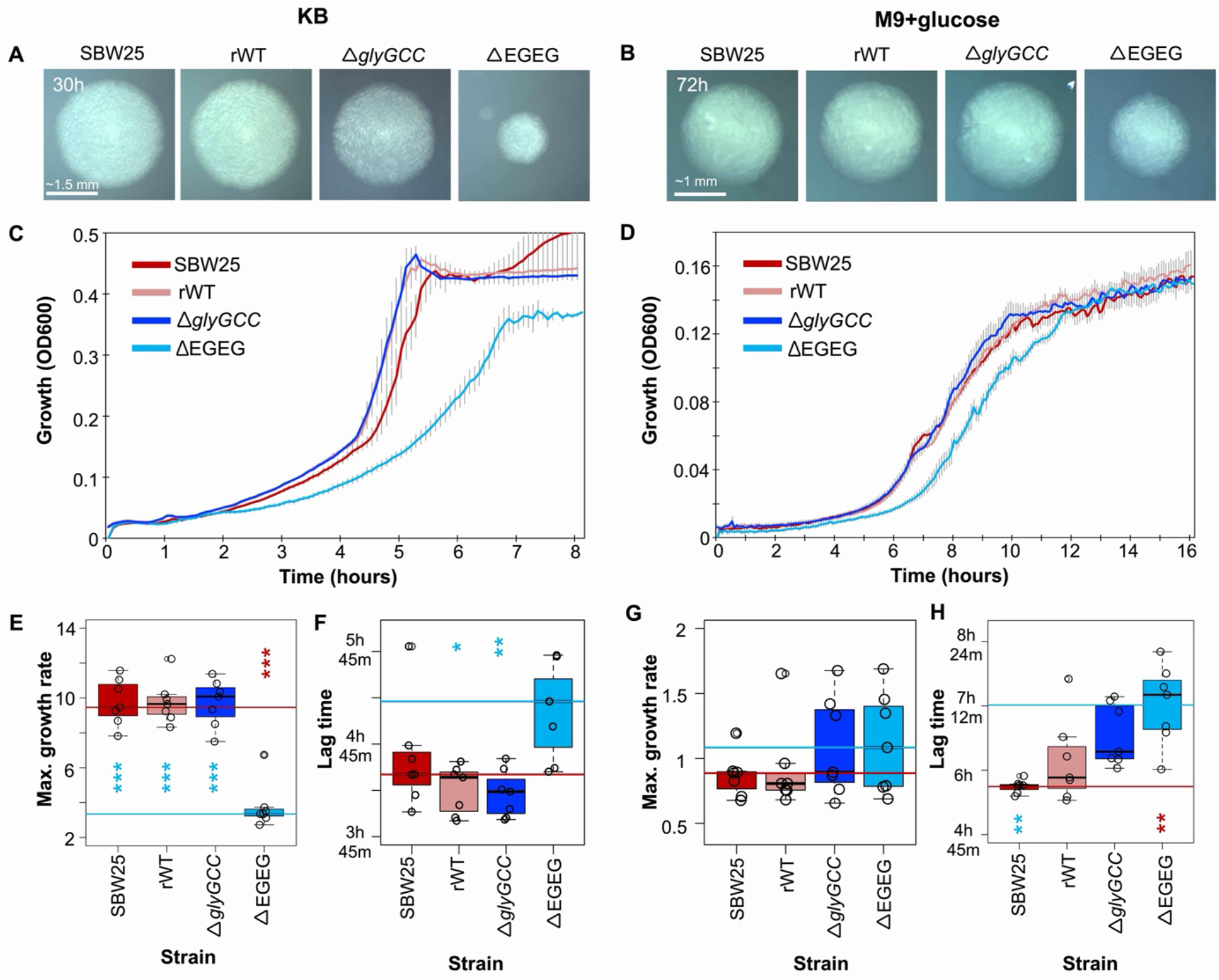
Deletion of the four gene tRNA locus *EGEG* reduces SBW25 growth in rich, undefined (KB) and poorer, defined (M9+glucose) media. (**A**-**B**) Colonies of SBW25, reconstructed wildtype (the engineering control, rWT), and the two engineered genotypes Δ*glyGCC* and ΔEGEG at room temperature for 30 hours on KB agar, or 72 hours on M9 agar. Scale bar applies to all four images in each panel. (**C**-**D**) Growth (measured as absorbance at 600 nm) of the same genotypes in liquid KB for 8 hours, or M9 for 16 hours. Lines show the mean of 7 replicates at 5-minute intervals; error bars are ± 1 SE. (**E-H**) Maximum growth rates (change in absorbance; mOD min^-1^) and lag times of growth profiles in panel C (KB), and panel D (M9). In all four panels, red and blue lines are the medians of SBW25 and ΔEGEG, respectively. Non-parametric Dunn’s tests, followed by the Benjamini-Hochberg procedure to correct for multiple comparisons, were used to test for a difference in medians between pairs of genotypes in each panel. Stars indicate statistically significant *p-*values for a difference from SBW25 (red stars) or ΔEGEG (blue stars).

### The ΔEGEG growth defect is rapidly compensated by evolution

In order to determine whether, and how, the ΔEGEG growth defect can be genetically compensated, a serial transfer evolution experiment was performed. Ten independent lines were passaged through repeated cycles of growth and dilution in shaken rich medium (KB), for 28 days (a total of ∼200 generations over 28 days; 1 % daily bottleneck). Five evolutionary lines were founded by ΔEGEG (lines M1-M5), and five by the wildtype (control lines W1-W5). Within 21 days (∼150 generations) each of the five mutant lines showed visibly improved growth, while the wildtype lines did not (from this point on, wildtype line W1 is used as a representative control line) (see Supplementary Figure S1). Single genotypes purified from day 21 of each mutant line (namely, genotypes M1-1, M2-1, M3-1, M4-1, and M5-1) showed improved growth on agar plates and in liquid culture, with the growth profile of one genotype (M3-1) approximating that of the wildtype (Figure 3A-D). A control genotype, W1-1, isolated from day 21 of the representative wildtype line showed a growth profile comparable to that of SBW25 (Figure 3A-D).

**Figure 3.**
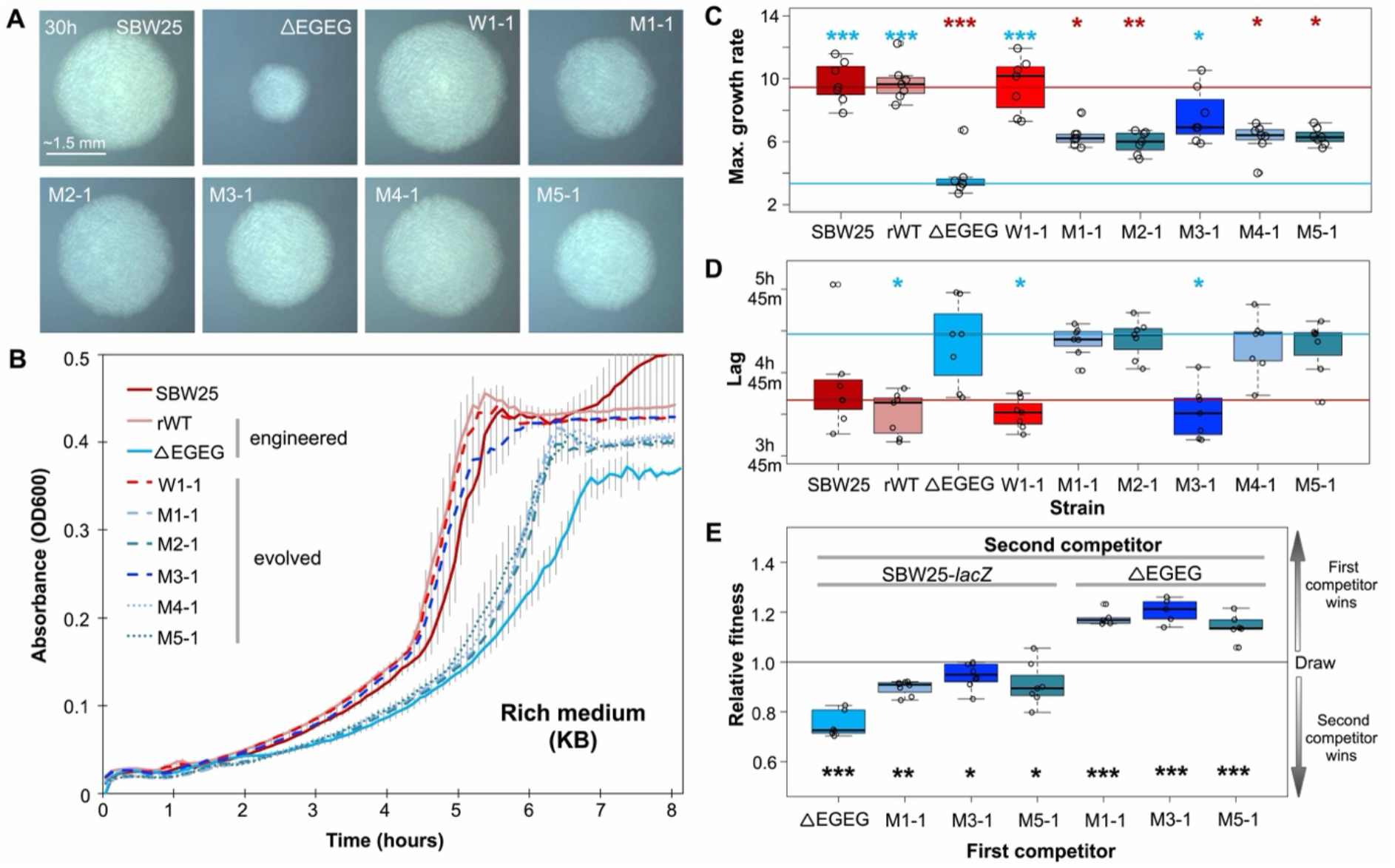
The growth defect caused by deleting four-gene tRNA locus *gluTTC-glyGCC-gluTTC-glyGCC* (*EGEG*) undergoes repeated compensation during a serial transfer experiment. (**A**) Colonies of founders SBW25, ΔEGEG, and six genotypes from day 21 of the serial transfer evolution experiment, grown on KB agar for 30 h at room temperature. Scale bar applies to all images. (**B**) Growth in liquid KB culture. Lines show the mean of 7 replicates at 5-minute intervals; error bars are ± 1 SE. (**C**,**D**) Maximum growth rate (change in absorbance; mOD min^-1^) and lag time for each growth profile in panel B. In both panels, red and blue lines are the medians of SBW25 and ΔEGEG, respectively. Non-parametric Dunn’s tests, followed by the Benjamini-Hochberg procedure to correct for multiple comparisons, were used to test for a difference in medians between pairs of genotypes. Stars indicate statistically significant *p-*values for a difference from SBW25 (red stars) or ΔEGEG (blue stars). The data for SBW25, rWT and ΔEGEG also appears in Figure 2. (**E**) Box plots of the relative fitness of competitor 1 (*x*-axis) and competitor 2 (horizontal bars at top). Direct, one-to-one competitions were performed in liquid KB for 24 hours (28°C, shaking). Between 5 and 8 replicate competitions were performed for each genotype pair. Relative fitness >1 means that the first competitor wins, <1 means that the second competitor wins. Statistically significant deviations of relative fitness from 1 were determined using either one-tailed one-sample *t*-tests, followed by a Benjamini-Hochberg correction for multiple comparisons. Data points from all replicates are overlaid on boxplots, and \*\*\**p*<0.001, \*\**p<*0.01, *p<0.05.

Three of the evolved mutant genotypes – M1-1, M3-1, and M5-1, each representing a different class of compensatory mutation (see next section) – were chosen for one-to-one competition experiments with (i) ΔEGEG, and (ii) a neutrally marked wildtype genotype (SBW25-*lacZ*). Each of the evolved mutant genotypes outperformed ΔEGEG (two-tailed one-sample *t*-tests *p*<0.0006), but were themselves outperformed by SBW25-*lacZ* (two-tailed one-sample *t*-tests *p*<0.05) (Figure 3E). Together, these results demonstrate that the growth defect caused by *EGEG* deletion can be readily compensated, at least in part, by evolution.

### Compensation of *EGEG* loss occurs via large-scale, tandem duplications spanning *glyGCC*

In order to identify the mutation(s) behind compensation, whole genome re-sequencing was performed on the six genotypes isolated from day 21 of the evolution experiment (M1-1, M2-1, M3-1, M4-1, M5-1, and W1-1). Per sample, over one million, 250-bp long Illumina sequencing reads were aligned to the SBW25 reference genome (8), resulting in a minimum mean coverage of 36.4 reads per genomic base. *Breseq* (64) and downstream analyses were used to predict the mutations in each evolved genotype (Supplementary Table S2).

In each of the five genotypes isolated from a mutant line, a large, direct, tandem duplication was identified at around ∼2.38 Mb of the SBW25 chromosome (Figure 4A-C). No evidence of any such duplication fragments was found in the representative wildtype control genotype, strongly suggesting that these large duplications play an important role in the observed compensatory effect. In one mutant genotype (M5-1), an additional, non-synonymous SNP was identified in *rplC* (located outside of the duplicated region; see Figure 4B). While the *rplC* mutation affects the translational apparatus (via amino acid change T43I in the 50S ribosomal protein L3), no evidence was found for a role in the compensated phenotype of M5-1: in the absence of the accompanying duplication, the *rplC* mutation did not improve the growth of ΔEGEG (see Supplementary Figure S2).

**Figure 4.**
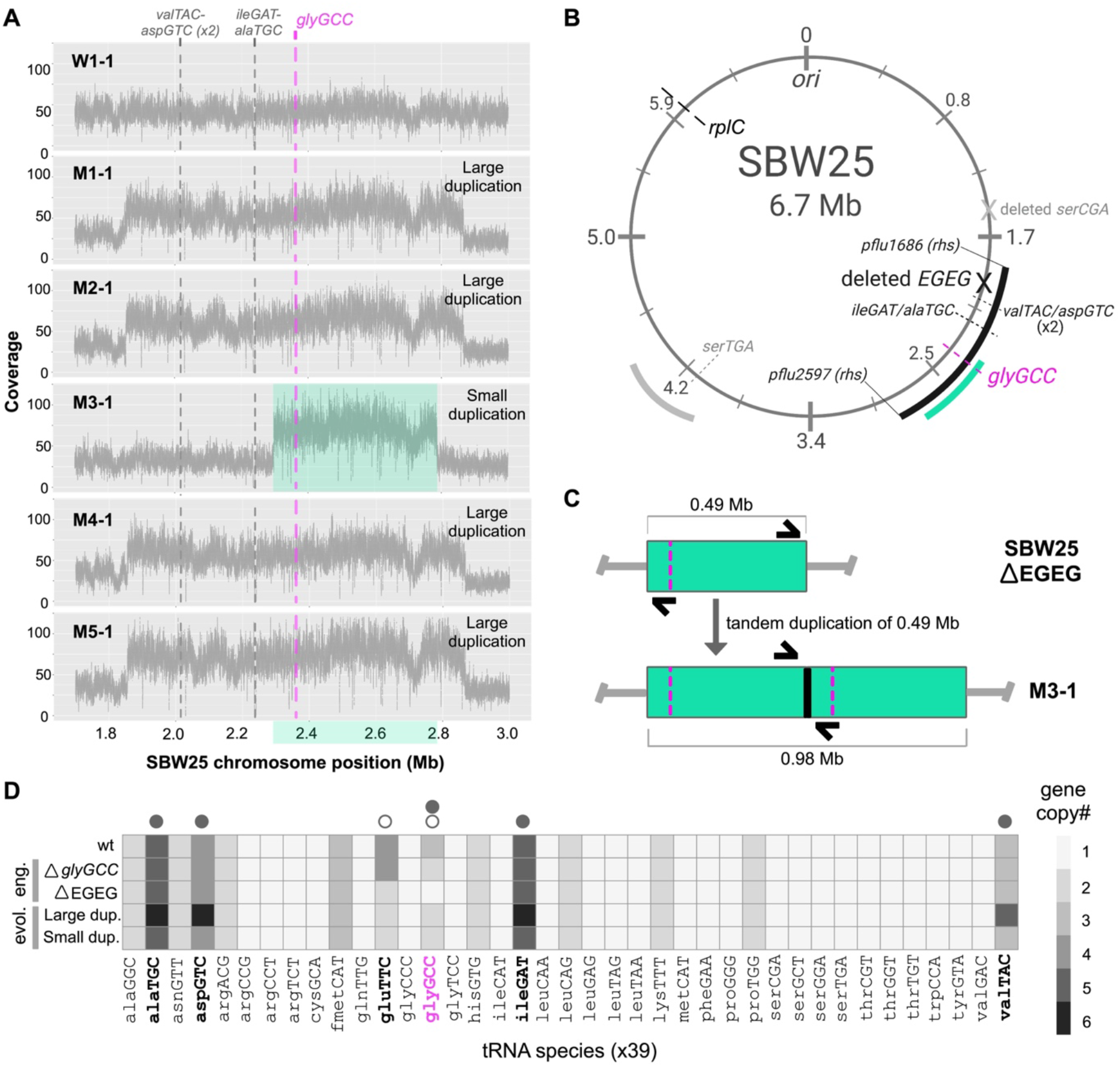
Large-scale, tandem duplications compensate for deletion of *EGEG*. **(A)** Number of raw sequencing reads covering 1.8 - 3.0 Mb of the reference genome. A jump to two-fold coverage is evident in the genotypes isolated from mutant populations (M1-1 to M5-1), and not in the control genotype isolated from the representative wildtype population (W1-1). The ∼0.49 Mb region of double coverage shared by all mutants is highlighted in green; tRNA genes are shown by dotted lines. (**B**) Cartoon depicting the positions of duplicated regions on the SBW25 genome. Black arc=∼1 Mb duplication (in M1-1, M2-1, M4-1, M5-1); green arc=∼0.49 Mb duplication (in M3-1); grey arc=distinct ∼0.4 Mb region in which duplication fragments have previously been reported (6). Duplicated tRNA genes are marked by dotted lines; deleted tRNA genes are marked with a cross. Image drawn using BioRender.com. (**C**) Depiction of the duplication event in M3-1, resulting in an additional ∼0.49 Mb of DNA and double copies of all genes in the duplicated region. Thick black line=duplication junction, which can be amplified using a duplication junction PCR (primer positions in black; see Supplementary Text S2). (**D**) Heatmap showing differences in the gene copy numbers coding for 39 tRNA species between wildtype (wt), engineered (eng.), and evolved (evol.) genotypes. Open/closed circles=tRNA species whose gene copy number changed by engineering/evolution.

The duplicated region in each ΔEGEG-derived genotype was more precisely defined using computational analyses and duplication junction PCRs (Table 1, Supplementary Text S2). While each duplication fragment has distinct endpoints, the fragments fall into two broad classes based on size: ∼1 Mb and ∼0.49 Mb duplications. ∼1 Mb duplications were identified in four evolved genotypes (M1-1, M2-1, M4-1, and M5-1). This duplication type typically occurs between long stretches of repetitive DNA at ∼1.86 Mb and ∼2.86 Mb of the SBW25 chromosome, and encompasses over 900 putative genes (Supplementary Table S3). The ∼0.49 Mb duplication was identified in one genotype, M3-1, between positions ∼2.30 Mb and ∼2.79 Mb of the SBW25 chromosome. At approximately half the size of the ∼1 Mb duplication, the ∼0.49 Mb duplication is predicted to contain only 426 genes (Supplementary Table S3). Notably, both types of duplication contain distinct subsets of tRNA genes: the ∼1 Mb fragments encompass seven tRNA genes (resulting in 69 tRNA genes in the evolved genome), while the ∼0.49 Mb fragment contains one tRNA gene (giving 63 tRNA genes) (Figure 4D).

**Table 1.** Details of duplications identified in five compensated genotypes isolated from the five evolving mutant populations on day 21 of the evolution experiment. Base positions refer to the SBW25 wildtype genome sequence (NCBI accession number: NC_012660.1). For a list of the genes contained within each duplication fragment, see Supplementary Table S3.

| Geno<br>type | Dup. size<br>(bp) | #<br>Dup.<br>genes | # Dup.<br>tRNA<br>genes | Junct.<br>name | Junction side 1 |  | Junction side 2 |  |
| --- | --- | --- | --- | --- | --- | --- | --- | --- |
|  |  |  |  |  | Base | Locus | Base | Locus |
| M1-1 | 1,016,166 | 917 | 69 | M1junct | 2,864,810 | *pflu2597*<br>(*rhs*) | 1,848,645 | Intergenic<br>(no repeat) |
| M2-1 | 1,012,338 | 911 | 69 | M2junct | 2,865,243–<br>2,865,277 | *pflu2597*<br>(*rhs*) | 1,852,904–<br>1,853,090 | *pflu1686*<br>(*rhs*) |
| M3-1 | 489,709 | 426 | 63 | M3junct | 2,786,207 | *pflu2547*<br>(no repeat) | 2,296,499 | Intergenic<br>(repeat) |
| M4-1 | ~1,012,000 | 911 | 69 | M4junct | ~2,865,500 | *pflu2597*<br>(*rhs*) | ~1,853,600 | *pflu1686*<br>(*rhs*) |
| M5-1 | 1,012,339 | 911 | 69 | M5junct | 2,866,737–<br>2,866,785 | *pflu2597*<br>(*rhs*) | 1,854,339–<br>1,854,447 | *pflu1686*<br>(*rhs*) |

Both ∼1 Mb and ∼0.49 Mb duplications encompass many hundreds of genes, with each constituent gene having a particular effect – positive, negative, or neutral – on growth. Since the net effect of each duplication fragment is compensatory, at least one gene in each fragment is presumed to have a sizeable positive effect on ΔEGEG growth. To gain insight into the gene(s) that may contribute to this positive effect, we focused on the ∼0.49 Mb fragment, for two reasons: (i) this region is duplicated in all five evolved genotypes, and (ii) it affords the greatest growth advantage of all observed duplication fragments. Of the 426 putative genes in the ∼0.49 Mb fragment, only one has an obvious role in translation: *glyGCC*, the sole remaining gene encoding tRNA-Gly(GCC) in ΔEGEG (see Figure 1). Interestingly, none of duplicated genes present a clear link to the co-deleted *gluTTC* genes. Hence, we hypothesize that changes in *glyGCC* gene copy number are the major determinant of the observed fitness effects.

### The observed tRNA gene set changes affect tRNA-Gly(GCC) levels

We sought to test whether the above *glyGCC* gene copy number changes are reflected in mature tRNA pools, using YAMAT-seq (a deep-sequencing technique that quantifies the relative proportions of tRNA species in the mature tRNA pool) (6, 67). YAMAT-seq was performed on 24 samples, including three replicates of each of eight genotypes: wildtype SBW25, the two engineered genotypes with reductions in *glyGCC* copy number (ΔEGEG, Δ*glyGCC*), three evolved mutant genotypes that reflect the identified compensatory mutations (M1-1, M3-1, and M5-1), and two control genotypes (engineering control rWT, evolution control W1-1). Illumina sequencing resulted in a minimum of 1.47 million raw tRNA reads per sample; these were aligned to a set of reference sequences, consisting of all the unique tRNA gene sequences predicted for SBW25 (6, 7) (see Supplementary Text S4). Each of the expected 39 tRNA species was detected in every sample, in varying proportions (Supplementary Table S4). Notably, tRNA-Gly(GCC) – encoded by three *glyGCC* genes – is the most highly represented of all tRNA species in the wildtype, accounting for 11.0 ± 1.0 % (mean ± SE) of the mature tRNA pool (Figure 5A). Four tRNA species (tRNA-Ile(CAU), tRNA-Phe(GAA), tRNA-Thr(CGU), and tRNA-Glu(UUC)) were detected only at very low levels (<0.1 % of the pool). It is probable that various post-transcriptional modifications present in these tRNA species reduced the efficacy of the YAMAT-seq protocol (specifically, the reverse transcription reaction; see Methods) (6, 67, 75). Due to unreliably low read numbers, these four tRNA species were removed from downstream analyses.

**Figure 5.**
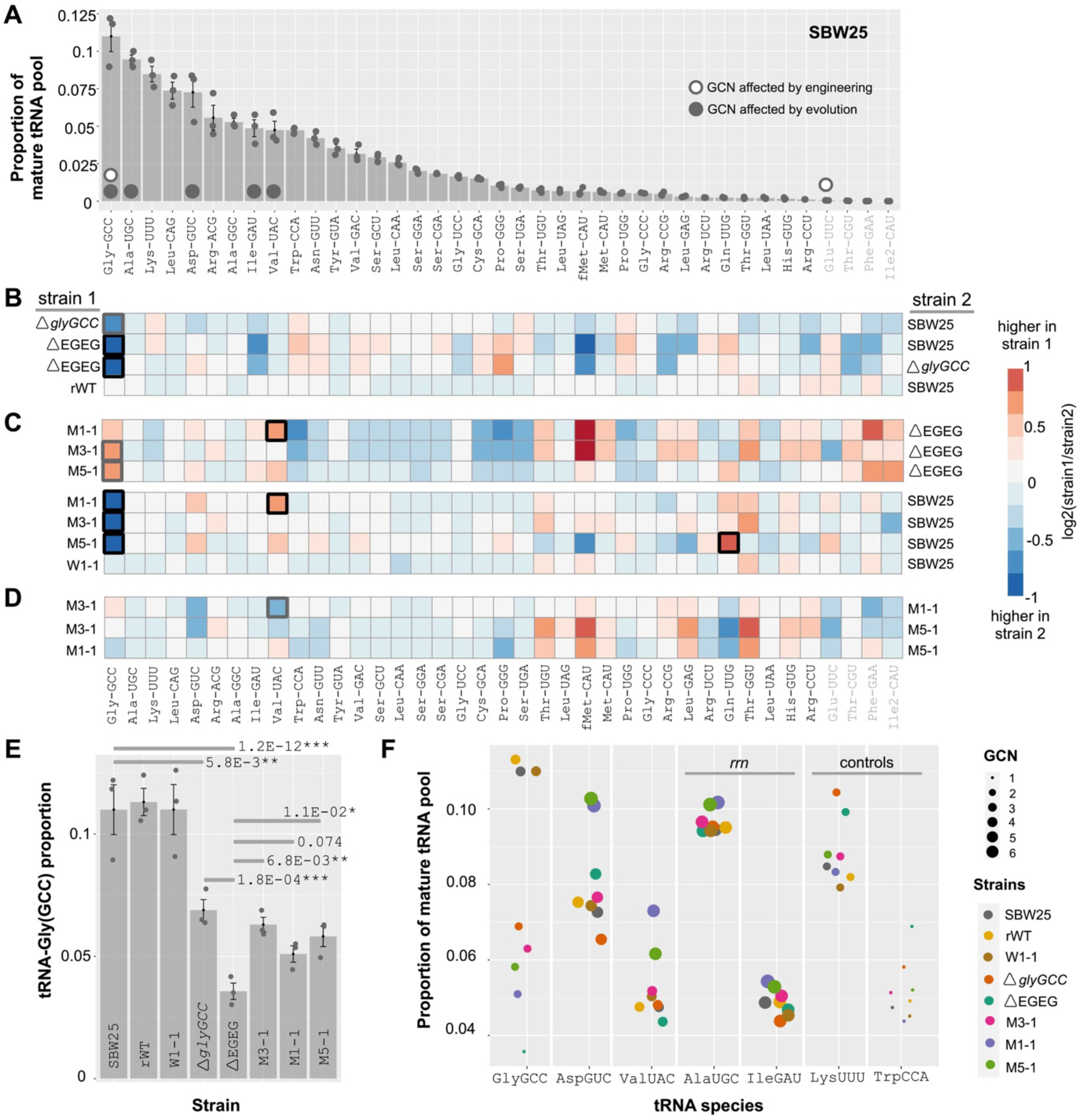
*glyGCC* gene copy number changes are mirrored in the mature tRNA pool. (**A**) Proportions of the 39 tRNA species in the SBW25 mature tRNA pool (highest to lowest). Light grey=excluded tRNA species. Open/closed circles=tRNA species whose gene copy number (GCN) changes by engineering/evolution in downstream experiments. (**B-D**) Heatmaps of differences in tRNA species proportions (expressed as log2-fold change(strain1/strain2); see Supplementary Table S5) between genotype pairs, to detect the effect of engineering (panel B) and evolution (panels C-D). Box outlines indicate statistical significance (DESeq2 adjusted p-values; thin line *p*>0.01; thick grey line 0.01 <*p*>0.001; thick black line *p*<0.001). (**E**) Bar graph of tRNA-Gly(GCC) proportion by genotype. Bars=means of 3 replicates ± 1 SE. DESeq2 adjusted *p*-values show tRNA-Gly(GCC) differences between genotypes pairs. (**F**) Scatter plot of mean proportions ± 1 SE of 7 tRNA species, by genotype. GCN for first 5 tRNA species varies by engineering and/or evolution (indicated by point size), while the final 2 are controls (constant GCN). *rrn*=tRNA species genes co-localize with rRNA operons.

DESeq2 analyses were used to detect differences in the relative proportions of all 39 tRNA species between genotype pairs of interest (Figure 5B-D, Supplementary Table S5) (6, 36, 69). Among the measurable tRNA species, the predominant – and in many cases, only statistically significant – effect of the engineering and evolution experiments was on tRNA-Gly(GCC) (Figure 5B-E). Deletion of either *glyGCC* or the *EGEG* locus saw a reduction in the relative contribution of tRNA-Gly(GCC) to the mature tRNA pool. Specifically, compared with 11.0 ± 1.0 % of the wildtype tRNA pool, tRNA-Gly(GCC) accounts for only 6.9 ± 0.43 % and 3.6 ± 0.34 % of the Δ*glyGCC* and ΔEGEG mature tRNA pools, respectively. The severe reduction in tRNA-Gly(GCC) in ΔEGEG was partially reversed by adaptive evolution; genotypes M1-1, M3-1, and M5-1 show tRNA-Gly(GCC) levels of 5.1 ± 0.34 %, 6.3 ± 0.30 %, and 5.8 ± 0.42 %, respectively. No such changes were observed in the tRNA pools of the engineering or evolution control genotypes (rWT, W1-1) (Figure 5B-E). Unfortunately, due to the unreliably low read numbers obtained, we are unable to comment on the effect of *EGEG* deletion, or subsequent evolution, on tRNA-Glu(UUC).

In addition to tRNA-Gly(GCC), the ∼1 Mb duplication carried by genotypes M1-1 and M5-1 changes the number of genes coding for a further four tRNA species: tRNA-Ala(UGC) (from five to six gene copies), tRNA-Asp(GUC) (from four to six gene copies), tRNA-Ile(GAU) (from five to six gene copies), and tRNA-Val(UAC) (from three to five gene copies). While the ∼1 Mb duplication fragment increases the contribution of each of these tRNA species to the mature tRNA pool, these changes are less pronounced than those observed for tRNA-Gly(GCC) (and very few are statistically significant) (Figures 5C, 5F). One reason for these weaker effects could be that each of the additional four tRNA species is encoded by multiple, dispersed tRNA gene copies, only some of which lie within the duplicated region. The degree to which each of these dispersed gene copies contributes to the final level of the mature tRNA species may vary considerably, and remains unknown (76, 77). Finally, we note that in addition to incurring no detectable growth or fitness effects (see Supplementary Figure S2), the *rplC* mutation had no major effects on the relative proportions of tRNA species in the mature tRNA pool, under the tested conditions; no statistically significant differences were detected between genotypes M1-1 (∼1 Mb duplication) and M5-1 (∼1 Mb duplication + *rplC* mutation) (Figure 5D).

The results in this section show that changes in *glyGCC* gene copy number are mirrored by changes in the proportional contribution of tRNA-Gly(GCC) to the mature tRNA pool. Specifically, reductions in *glyGCC* gene copy number result in progressively lower proportions of tRNA-Gly(GCC), which we hypothesize eventually manifest as a reduction in growth rate. Growth rate is subsequently restored by large-scale, tandem duplication events, one effect of which is to elevate *glyGCC* gene copy number and (partially) reinstate tRNA-Gly(GCC) levels.

### Duplication fragments are unstable, leading to genetically mixed populations

Large-scale, tandem duplications are typically highly unstable; they are known to be lost at high rates via homologous recombination events during DNA replication (often leaving no trace in the segregant genome) (55, 78, 79). The duplication fragments observed in this work are no exception. When grown in overnight culture and plated on agar plates, each of the duplication-carrying genotypes repeatedly gives rise to colonies of two distinct types: (i) a majority of larger, wildtype-like colonies that retain the duplication fragment, and (ii) a sizeable minority of smaller, ΔEGEG-like colonies in which the initial duplication fragment has been lost (Figure 6A-B; see also Supplementary Figure S2).

**Figure 6.**
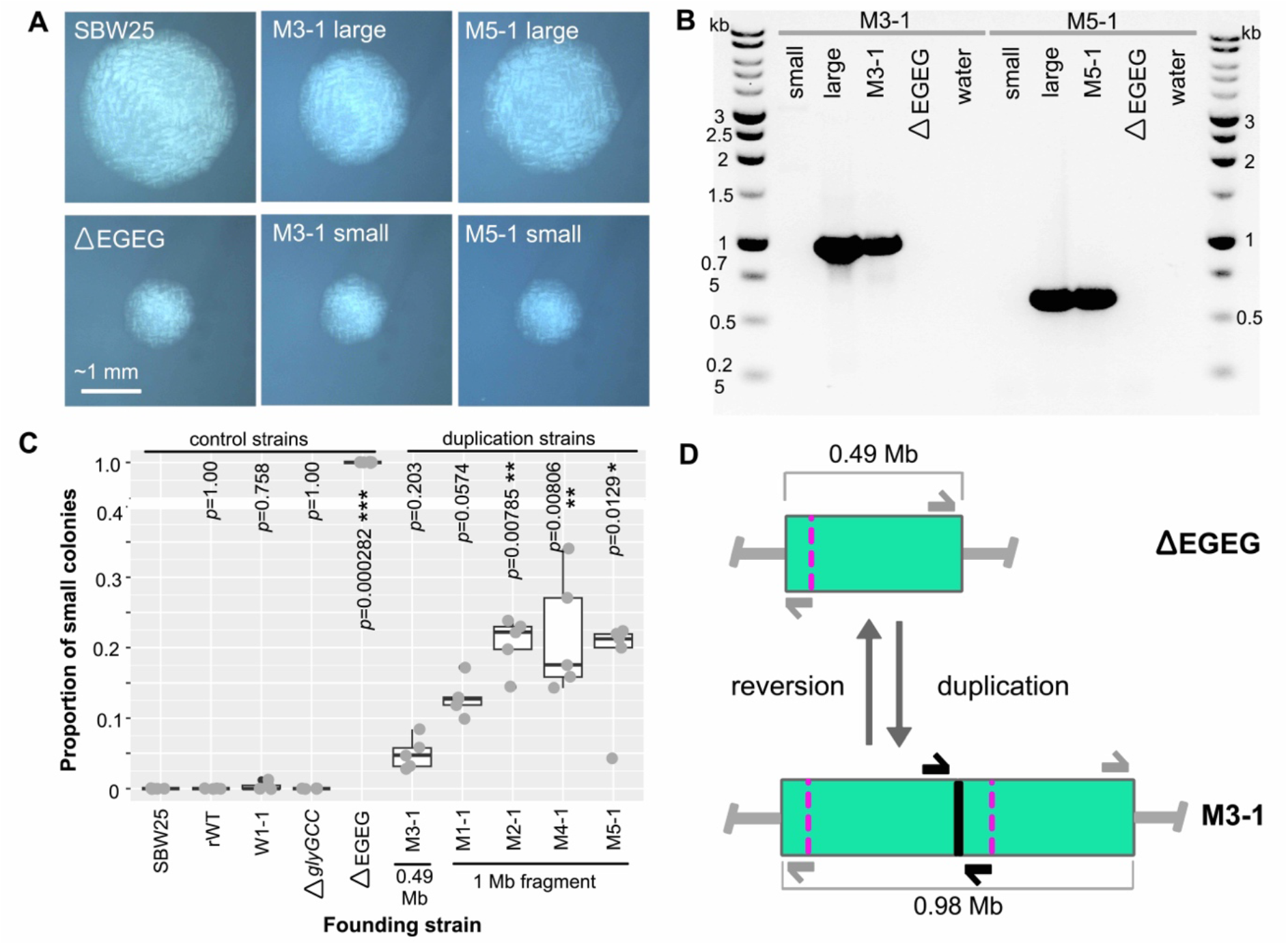
The large-scale, tandem duplications are rapidly lost in overnight culture. (**A**) Two colony morphologies – large and small – are observed when plating from overnight cultures of duplication-carrying genotypes (colonies from duplication genotypes M3-1 and M5-1 are presented here). All images taken under the same magnification after ∼48 hours’ incubation on KB agar at room temperature (∼21°C). Scale bar applies to all images in panel. (**B**) Small colonies no longer amplify the unique junction introduced by the relevant duplication fragment (illustrated by the thick black lines in M3-1 cartoon of panel D). (**C**) Duplication fragment loss was quantified as the proportion of small colonies when plating from an overnight culture. Five replicates were included per genotype, and at least 41 colonies were counted per replicate. Boxplots summarize the data (medians), and grey dots are individual replicates. Statistical significance of a difference in the median proportion of small colonies compared to SBW25 (wildtype) was calculated using a Dunn test (with Benjamini-Hochberg correction for multiple comparisons). Significance levels: no stars (*p*>0.05), *0.05<*p*<0.001, **0.01<*p*<0.001, \*\*\**p*<0.001. (**D**) Cartoon depicting the gain and loss of duplication fragments, leading to a mixed population of the duplication genotype (here, M3-1) and ΔEGEG.

The degree of duplication fragment loss was quantified using a stability test, in which the proportion of small colonies (*i.e.*, segregants) arising from independent cultures founded by each of the five duplication-carrying, evolved mutant genotypes was determined. Between 3 % and 34 % of colonies arising from duplication genotype cultures showed the small, ΔEGEG-like phenotype (Figure 6C). The proportion of small colonies observed was dependent on the size of the initial duplication fragment, with small and large duplication fragments giving rise to respective averages of 5.0 % ± 1.0% and 18 % ± 1.5 % segregants (mean ± SE; non-parametric two-sample *t*-test for difference between means *p*=3.5e-07***). The higher proportion of segregants in populations founded by duplication genotypes carrying the larger, ∼1 Mb fragments presumably results from a combination of the higher number of recombination events occurring across a more extensive region of homology, and the lesser fitness advantage afforded by the ∼1 Mb duplications (see Figure 3).

The results in this section demonstrate that the adaptive duplication events are inherently unstable; they, along with their compensatory tRNA gene copies, are lost at high rates. In growing populations founded by duplication-carrying genotypes, this high rate of loss leads to the presence of at least two genetically distinct sub-populations of cells: (i) those carrying the initial duplication fragment, and (ii) those in which the fragment has been lost (*i.e.*, ΔEGEG) (Figure 6D).

### Multiple duplication fragments rapidly arise and compete within evolving populations

The ease with which duplications arise and are lost suggests complex evolutionary dynamics within the mutant lines. To begin investigating, we tracked ∼1 Mb and ∼0.49 Mb duplication fragments through one of the evolutionary lines in which they were identified by the day 21 genotype sequencing (M2 and M3, respectively). This was done by normalizing the intensity of a duplication junction PCR band (which is expected to change as the population evolves) with that of a product amplified from a genomic region unaffected by the duplication events (which is expected to remain constant) (see Methods, Supplementary Figure S3, and Supplementary Table S6). Across an evolving population, changes in the frequency of the target duplications are expected to be reflected by changes in the normalized intensity values (we note that the normalized intensities are not intended to provide precise frequency estimates, but to reveal patterns of change).

The normalized intensity profile of the ∼1 Mb fragment across line M2 is shown in Figure 7A, and that of the ∼0.49 Mb fragment across line M3 is in Figure 7B. Both duplications were detected early in their respective lines (∼days 4-6), and spread rapidly to peak frequency (∼days 12-18). Thereafter, the ∼1Mb fragment steadily declined to low levels, while the smaller (and hence fitter and more stable) ∼0.49 Mb fragment was maintained at high levels until the end of the experiment.

**Figure 7.**
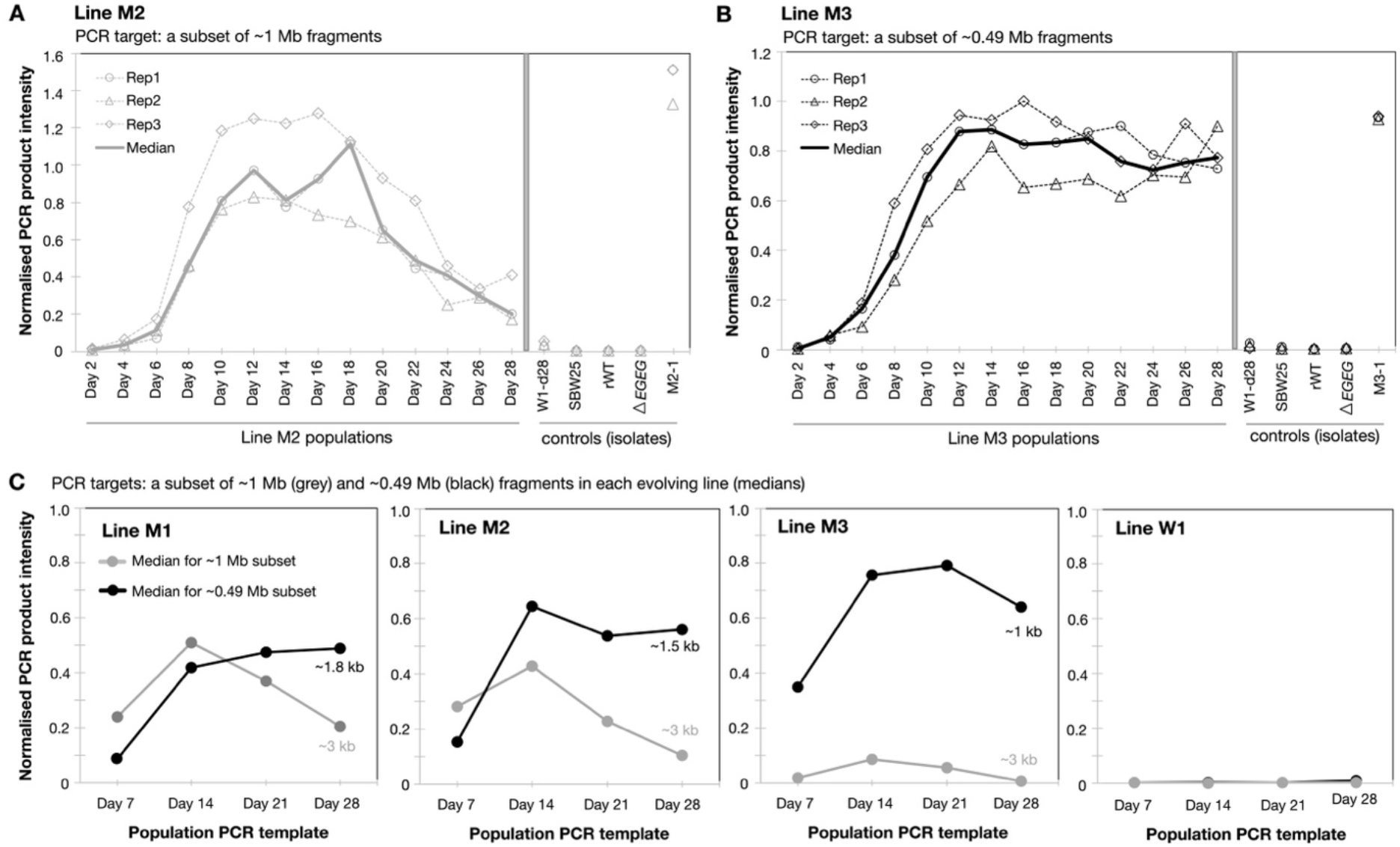
The complex evolutionary dynamics of large-scale duplication fragments in evolving mutant populations. Duplication junction PCRs were used to follow the dynamics of subsets of ∼1 Mb and ∼0.49 Mb duplications across evolving populations. The intensity of the PCR product obtained across each emergent duplication junction was measured relative to a PCR product amplified from a control region located outside of the genomic region affected by duplication events. Various control samples – in which (almost) all or none of the population is expected to carry duplication fragments – were included in each set of PCRs (see Methods for further details). The subsets of ∼1 Mb (**A**; primer pair M1and2_junct_f/M5_junct_r) and ∼0.49 Mb (**B**; primer pair M3_junct_f/r) duplications show different evolutionary trajectories in the lines in which they were originally identified; the ∼1 Mb fragment rises and then wanes in line M2, while the ∼0.49 Mb fragment rises to, and is maintained for longer at, a higher level in line M3. (**C**) Multiple duplication fragments arise and compete within evolving populations. Each PCR was performed in triplicate, and the median is presented here (for full data sets see Supplementary Figure S5A; Supplementary Table S6).

The above results hint at the presence of multiple adaptive genotypes in at least some evolving populations. To further explore this possibility, normalized duplication junction PCRs were used to test for the co-existence of ∼1 Mb and ∼0.49 Mb duplication fragments in evolving lines M1, M2, M3, as well as a representative control line (W1). (Lines M4 and M5 were excluded due to the discovery of external contaminants in the later stages of the experiment, see Supplementary Figure S1.) The results presented in Figure 7C show that both the ∼1 Mb and ∼0.49 Mb duplications are indeed detectable in all three evolving mutant lines (and not in W1). The co-existence of both fragment types is most prominent during the early stages of the experiment (∼days 7-21); in general, the ∼1 Mb fragment declines in the later stages of the experiment, while the ∼0.49 Mb fragment remains at relatively constant levels. Notably, distinct subtypes of the smaller duplication fragment – each with subtly different endpoints and hence variably sized duplication junction PCR products – were detected in lines M1-M3 (see Supplementary Figures S4 and S5A for more detail). Further, since each duplication junction PCR tests only for a subset of hypothetically possible duplication fragments, we note that there may be other duplications present that are not captured by these PCRs.

The results in this section provide evidence of complex evolutionary dynamics of duplications in the evolving mutant populations; both ∼1 Mb and ∼0.49 Mb fragments, each typically occurring between sets of long repeats, arise readily and repeatedly within evolving ΔEGEG populations. Over time, the larger ∼1 Mb fragments are outcompeted by their smaller (and fitter/more stable) ∼0.49 Mb counterparts.

### Smaller duplication fragments arise and displace larger duplication fragments

To further investigate the evolutionary dynamics of duplications, we performed population-level whole genome re-sequencing on the day 28 populations of mutant lines M1-M3 (day 28 wildtype populations W1-W5, and an ancestral SBW25 sample were included as controls). For each of the nine samples, 150-bp paired-end Illumina reads were aligned to the SBW25 reference sequence (8) using *breseq* (on polymorphism detection settings, see Methods) (64), giving a minimum mean coverage of 154 reads per genomic base. The raw *breseq* output, and a list of curated mutation predictions, are provided in Supplementary Table S7.

From the *breseq* output and further computational analyses (see Supplementary Text S3), eleven distinct duplication fragments were identified across the three day-28 ΔEGEG-derived populations (Table 2). No such duplications were observed in any of the control samples. Ten of these eleven duplication fragments are novel; the eleventh is identical to the ∼0.49 Mb fragment previously identified in the genotype M3-1 (isolated from the day-21, line M3 population). Each of the three day-28 mutant populations was found to contain at least three distinct, co-existing duplication fragments, including multiple (relatively) large fragments (between ∼0.28 Mb and ∼0.77 Mb in size, often with endpoints in repetitive DNA regions), and one much smaller fragment (between ∼2.2 kb and ∼4.3 kb, with no obvious endpoint homology) (Figure 8A). Notably, each mutant population is predicted to contain a ∼0.49 Mb duplication fragment between genomic positions ∼2.30 Mb and ∼2.79 Mb (*i.e.*, is expected to give a PCR product with the duplication junction primers used in Figure 7C).

**Figure 8.**
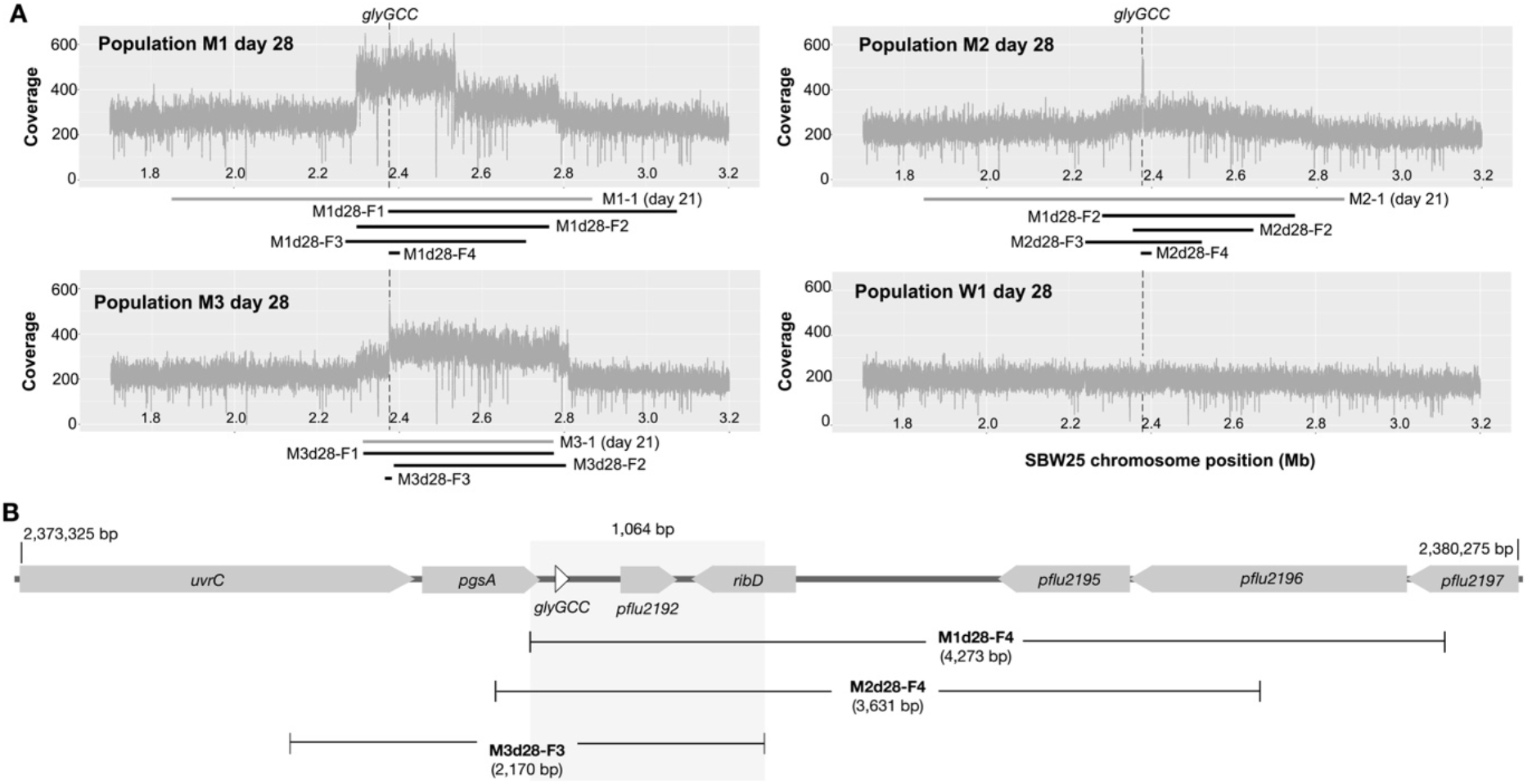
Eleven distinct duplication fragments are detected in mutant lines M1, M2, and M3 on day 28 of the evolution experiment. (**A**) Number of raw sequencing reads covering 1.7 Mb – 3.2 Mb of the reference genome, shown for the three mutant lines (M1, M2, M3) and one representative wildtype control line (W1) on day 28. Large changes in coverage can be seen in all mutant populations (but not the wildtype population). The black bars below each plot indicate approximate positions of each computationally predicted duplication (see Table 2); grey bars show the duplication identified in the corresponding day 21 isolate (see Table 1). (**B**) Structure of the ∼7 kb genomic region containing the three smallest duplication fragments. Grey highlighting indicates the 1,064 bp that is duplicated in all three fragments. This region contains two complete genes: tRNA gene *glyGCC* and predicted pseudogene *pflu2192* (see Supplementary Table S3).

**Table 2.**
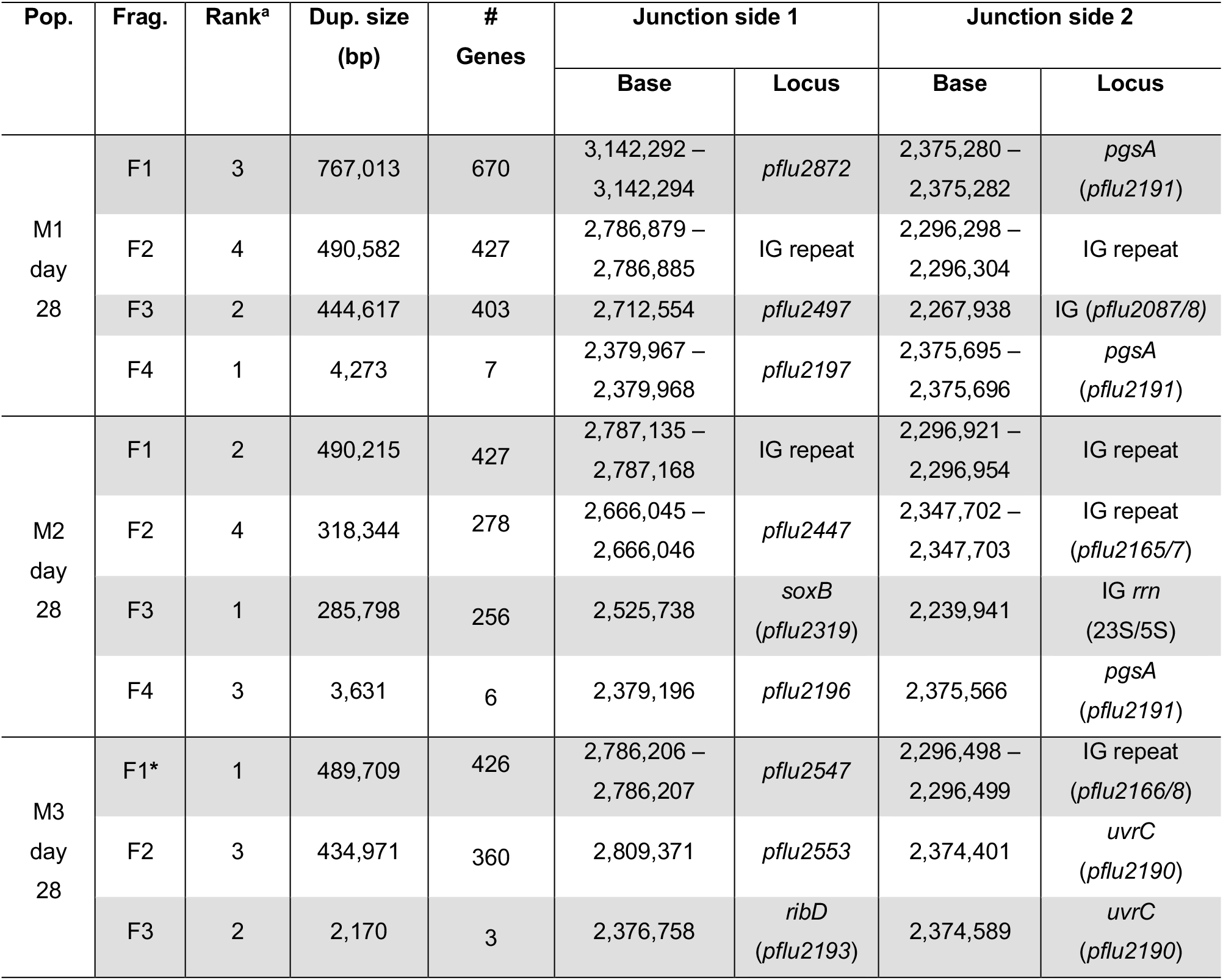
Details of the duplications computationally predicted in day 28 populations from evolving mutant lines M1, M2, and M3. Base positions refer to the SBW25 wildtype genome sequence (NCBI accession number: NC_012660.1). For a list of the genes contained within each predicted duplication, see Supplementary Table S3. ^a^Duplications are ranked within each population in descending order of prevalence (according to the adjusted number of Illumina reads perfectly matching the complete proposed duplication junction sequence; see Supplementary Text S3). **\***Fragment M3d28-F1 is identical to the duplication identified in day 21 genotype M3-1 (see Table 1).

The smallest duplication fragment is predicted in line M3. It is just 2,170-bp long and contains only three complete genes: *pgsA* (predicted to be involved in phospholipid biosynthesis)*, glyGCC* (tRNA gene), and *pflu2192* (a putative pseudogene of phage origin) (Figure 8B). When all 15 of the duplication fragments identified in this study are considered, only two genes are duplicated in every fragment: *glyGCC* and *pflu2192* (Figure 8B, Supplementary Table S3). Taken together, the population sequencing and duplication junction PCRs provide strong support for the hypothesis that the major compensatory element of the duplication fragments is *glyGCC*; over time, progressively smaller duplication fragments encompassing *glyGCC* (and pseudogene *pflu2192)* arise and spread in the evolving, ΔEGEG-derived populations.

## DISCUSSION

Various studies have reported adaptive changes in tRNA gene copy number in laboratory populations of bacteria, in response to changing translational needs (6, 40, 41). In these studies, changes in gene copy number are underpinned by large-scale, tandem duplications (6, 41) or amplifications (40). Such mutations are both pervasive and, often, mechanistically reversible: large-scale duplications typically arise, and are lost, extremely rapidly (43, 44, 78, 80, 81), in a manner that is reminiscent of contingency loci in pathogenic bacteria (82, 83). Their transient nature means that their contributions to population diversity, and downstream effects on evolution, can be underappreciated (44). In this work, we (i) demonstrate the contribution of large-scale duplications to population-level diversity in tRNA gene set content, and (ii) show that such duplications may provide the raw material for longer-term evolutionary changes in basal tRNA gene set content.

### Spontaneous duplications rapidly generate population-level tRNA set diversity

As outlined in the introduction, large-scale duplications typically arise spontaneously (during DNA replication), at high rates, in a wide range of organisms (43, 46, 54–57). For instance, it has been reported that – without invoking any growth advantage – around 10 % of cells in an overnight *Salmonella* culture carry a large duplication of some region of the chromosome, with between 0.005 % and 3 % of cells carrying a duplication of a given locus (42, 43, 55). In short, a growing population of initially isogenic cells is expected to rapidly consist of a mixture of genotypes that vary with respect to the copy numbers of many genes (including tRNA genes).

We have demonstrated that similar large-scale, tandem duplications readily occur in at least two regions of the *P. fluorescens* SBW25 chromosome: either side of the putative replication terminus, centered around genomic positions ∼2.38 Mb (this work) and ∼4.16 Mb (6) (see Figure 4B). In this work we have reported 15 distinct duplications at position ∼2.38 Mb, including five duplications identified in genotypes isolated from day 21 of the evolution experiment, and a further ten duplications predicted in the day 28 population sequencing (see Tables 1-2 and Supplementary Table S3). These 15 duplications range from ∼2.2 kb to ∼1 Mb in size, and each one increases the initial tRNA gene set (of ΔEGEG) from 62 tRNA genes to either 63 or 69 tRNA genes (see Figure 4D). In our earlier work, we observed five duplications at position ∼4.16 Mb (6). These five duplications are up to ∼290 kb, and each adds one tRNA gene copy to the initial genome (Δ*serCGA*) (6). The tRNA gene set changes incurred by large-scale duplications have phenotypic consequences: the additional tRNA gene copies generally lead to higher proportional contributions of the corresponding tRNA species to the mature tRNA pool (see Figure 5 and (6)). Taken together, these results show that large-scale duplications readily occur at multiple locations in the SBW25 chromosome, thereby providing a mechanism for the rapid generation of population-level diversity in mature tRNA pool content.

The duplications reported in this work are detected at unequal rates. For instance, examples of similar ∼1 Mb and ∼0.49 Mb duplication fragments are found in the early stages of each evolving mutant line (see Table 2, Figure 7). This observation is consistent with literature showing that some chromosomal regions – particularly those delineated by long stretches of homologous DNA (*e.g., rrn* operons) – give rise to duplication events at especially high rates (79). Indeed, the prevalent ∼1 Mb and ∼0.49 Mb fragment classes are both delineated by long stretches of homologous DNA: the ∼1 Mb duplication fragments are typically flanked by two *rhs* genes at positions ∼1.86 Mb and ∼2.86 Mb (∼3 kb of homologous sequence), while the ∼0.49 Mb fragments are generally delineated by intergenic repeat regions at ∼2.30 Mb and ∼2.79 Mb (∼1.1 kb of homology).

In addition to the above mechanisms, tandem duplications can arise – via alternative mechanisms, and at much lower rates – between regions with little or no DNA homology (79). Such alternative mechanisms presumably contribute to the generation of the smaller duplication fragments detected at the later stages of evolution in this work; these smaller fragments, many of which have no obvious endpoint homology, could conceivably arise at low rates either directly from the founder genotype (ΔEGEG), or from duplication-carrying genotypes (through the partial segregation of a duplication fragment). Indeed, an example of the latter was recently reported in an experiment examining the fate of an advantageous, large-scale (∼1.66 Mb), tandem duplication fragment in *Salmonella*; during 2,000 generations of positive selection, the initial duplication fragment repeatedly shrank to ∼0.66 Mb, via the loss of an internal ∼1 Mb segment (84, 85).

### Unstable but adaptive duplications as evolutionary stepping stones

Generally speaking, large-scale, tandem duplication fragments are inherently unstable; they are prone to loss through RecA-mediated recombination during DNA replication (55, 79). The duplication fragments described in this work are similarly precarious – the stability test showed that, despite the net fitness advantage that they provide, duplications are lost by between 3 and 34 % of cells in overnight culture (see Figure 6). The rate of segregation was higher for genotypes carrying the larger (∼1 Mb) versus the smaller (∼0.49 Mb) duplications (median loss rates of 19 % and 5 %, respectively). This difference presumably results from a combination of the larger duplications (i) carrying twice the length of homologous DNA for recombination (*i.e.*, the duplication fragment), and (ii) conferring a lesser fitness advantage (see Figure 3) (44).

The inherent instability of the large-scale duplications renders them unlikely to represent a long-term evolutionary solution. That is, over time, inherently unstable duplication fragments are expected to be displaced by a more stable type of mutation (*e.g.*, SNPs affecting tRNA gene expression, or smaller/more stable duplication fragments). Indeed, we already see evidence of such displacement over the course of our 28-day evolution experiment; the large, ∼1 Mb fragments detected at the early stages of each mutant line tested (M1-M3) falls to very low or undetectable levels by the end of the experiment (see the population-level duplication junction PCRs in Figure 7). Furthermore, in one mutant line (M3), the co-existing smaller/fitter ∼0.49 Mb fragment also appears to drop in frequency towards the end of the experiment, hinting at the presence of at least one additional, higher-effect mutation in this population. Indeed, subsequent population-level genome re-sequencing detected the presence of much smaller (and, presumably, fitter and mechanistically more stable) duplication fragments in the later stages of all three mutant lines; each day 28 mutant population contains at least one duplication fragment of <5 kb (see Table 2, Figure 8).

The small (<5 kb) duplications fragments provide considerable insight into the mechanism behind the compensatory effects of the duplications. Together, the three smallest fragments reduce the core, overlapping region of all 15 reported duplication fragments to just 1,064 bp, which encompasses only two complete genes: tRNA gene *glyGCC*, and predicted pseudogene *pflu2192* (see Figure 8B). This observation strongly supports the hypotheses that (i) the ΔEGEG growth defect is primarily caused by a drop in tRNA-Gly(GCC), and (ii) this defect is readily compensated by duplicating *glyGCC* and thus (partially) restoring tRNA-Gly(GCC) levels. We note that this does not rule out a more minor role for tRNA-Glu(UUC), the second tRNA species targeted by the initial *EGEG* deletion. For instance, it is conceivable that a secondary, lower-effect wave of adaptation could target *gluTTC*. Alternatively, selection pressure on *gluTTC* may be stronger under different experimental conditions.

The detection of progressively shorter duplication fragments raises the interesting question of what would occur if the evolution experiment were continued. One possibility is that the duplications would continue to reduce in size. Presumably, the smallest possible adaptive duplication is approximately ∼150-200 bp long (encompassing merely the *glyGCC* gene and promoter). Mechanistically speaking, such a short duplication fragment would be expected to be relatively stable, and hence a genotype carrying such a duplication could be considered to encode a new, comparatively stable tRNA gene set. The long-term evolutionary fate of the duplication fragments – and hence the new tRNA gene copies – remains to be seen, and will provide insight into the extent to which within-genome duplication events contribute to the evolution of extant bacterial tRNA gene sets.

## DATA AVAILABILITY

Whole genome sequencing data has been deposited with NCBI Short Read Archive (SRA) under accession numbers PRJNA790786 (isolate sequencing) and PRJNA1035442 (population sequencing).

YAMAT-seq data has been deposited with NCBI Gene Expression Omnibus (GEO) under accession number GSE196705.

The remaining raw data relating to this project have been deposited on Zenodo (doi: 10.5281/zenodo.7276660).

## SUPPLEMENTARY DATA

Supplementary Data are available at NAR online.

## ACKNOWLEDGEMENTS

The authors thank Frederic Bertels for useful comments on the manuscript.

## AUTHOR CONTRIBUTIONS

Multi-author contributions are listed alphabetically. Conceived and designed the experiments: GBA, JG, PN, SL, WYN, ZK. Performed the experiments: all authors. Analyzed the data: all authors. Wrote the paper: JG. Commented on the paper: all authors. Conceived the research: JG.

