## Supplementary Information for "Large-scale duplication events underpin population-level flexibility in tRNA gene copy number in *Pseudomonas fluorescens* SBW25"

The supplementary information contains the following:

Supplementary Texts S1-S4 (pages 2-16)
Supplementary Figures S1-S5 (page 17-22)
Supplementary Tables S1-S7 (legends for accompanying excel files; pages 23-24)
Supplementary References (page 25)

#### Supplementary Text S1

#### Extended experimental procedures: construction of engineered genotypes

##### *1.1 Deletion of the* EGEG *tRNA locus from SBW25 (construction of ΔEGEG)*

In engineered mutant ΔEGEG, a 420-bp stretch of DNA encompassing the four-gene *gluTTC-glyGCC-gluTTC-glyGCC* locus (*EGEG* for short) are removed from the wildtype SBW25 genome. The construction process used is scar-free (*i.e.*, no other trace of the construction process, such as antibiotic markers, are left in the genome of the deletion mutant).

Outline of the deletion process:

1. Primers delEGEG_1, 2, 3, and 4 were used to construct a 1,362-bp deletion fragment via SOE-PCR (Ho et al. 1989). The fragment was ligated into the pCR™8/GW/TOPO® vector and TA Cloning® kit (Invitrogen™), and the sequence checked by Sanger sequencing with commercial primers GW1 and/or GW2.
2. The confirmed deletion fragment was transferred from pCR™8/GW/TOPO® into the pUIC3 vector (Rainey 1999), using the *Bgl*II restriction sites incorporated into the deletion fragment construction primers.
3. The deletion fragment in the pUIC3 vector was exchanged with the homologous chromosomal fragment using scar-free, two-step allelic exchange (described in detail in (Zhang and Rainey 2007; Ayan et al. 2020)). The sequence of the manipulated locus in the resulting genotype, ΔEGEG, was confirmed by PCR amplification and Sanger sequencing with delEGEG_f/r.

| **Primer Name** | **Sequence (5'→3')[a]** | **Use** |
|---|---|---|
| delEGEG_1 | gaagatctGTTGACGTTGAGTTCCAGTCGGGTAAG | Deletion fragment construction |
| delEGEG_2 | CACGTACAACATTCACTCCCTACCG | Deletion fragment construction |
| delEGEG_3 | cggtagggagtgaatgttgtacgtgCAACGAAAACGCCGATCAGTGATGATC | Deletion fragment construction |
| delEGEG_4 | gaagatctGATTACCTGGGAGGCAAACTGGGTATTG | Deletion fragment construction |
| delEGEG_f | GTGACCTGACCATGTAGGGCTTCGAG | Final PCR & sequencing |
| delEGEG_r | GCAAGGAAGTCCAACTGACCGTTAC | Final PCR & sequencing |
| delEGEG_f2 | CGTCCGTTACATGGATGCAC | Final sequencing |
| delEGEG_r2 | CAATGGCGTGCAGTTCGG | Final sequencing |

[a]underline denotes *Bgl*II restriction sites (these are used for the cloning process, and are not included in the final, scar-free deletion). Capital letters denote the region annealing to the SBW25 genome sequence, and small letters denote non-annealing primer sequence. Highlighting relates to the nucleotide sequences below. Primers are also listed in Supplementary Table S1.

```
>wildtype (shown here: bases 2027887-2029844)
…gtgacctgaccatgtagggcttcgagcaggaagcgcacgtccttggccttggcttcggtctggatcggcgcggg
tgggttgttcaccgggatcgattcgccctgagccaccagcgtattgagttcggcgatgcgggccttgatggggc
gctggtcttgttgaggttcaacaggcgctggttgacgttgagttccagtcgggtaagcaacaacttgagcgccgg
ggacatcaactggccggggaaggtctcggtgatttcaccgcaacggcgcaggcgatcgttaagctcgatctcggt
gcgcttcttttccagcttgttgttttccaccacgtggttcatatagccaatggcgatcagtattacgatcccggc
tattaccagcagggtgatcatgagtggtgtcaccggtgtgacctctttatagggtttacagtggagtgtagtgac
tgggccatccggcggatagggctgatcgaccagttgcccaggtagtttgcttctctatagataggcgtccgttac
atggatgcacattgccatcatttgtcggagcgaactatagcgccttgtcggacgccagaatataggcgccaaagc
ctgggcgggcacaatcctcccataagcccggaaagcagggcgaagtcattgatttaaataaatttatccttgggg
gttgacgacctttcaatccatccatagaatgcgcgccacttacagcgtaaagcacacagcgaaacgcggtaggga
gtgaatgttgtacgtgtgtccccttcgtctagtggcctaggacaccgccctttcacggcggtaacaggggttcga
gtcccctaggggacgccaatatgcgggaatagctcagttggtagagcacgaccttgccaaggtcggggtcgcgag
ttcgagtctcgtttcccgctccaaatttaagcggtattgccttcgggtggtactgagtgaaaccagggcataatc
ttcggatttaatctctggacactgaaatacacaccatgtgtttcagttgtgtgtccccttcgtctagtggcctag
gacaccgccctttcacggcggtaacaggggttcgagtcccctaggggacgccatttgcgggaatagctcagttgg
tagagcacgaccttgccaaggtcggggtcgcgagttcgagtctcgtttcccgctccatattcaacgaaaacgccg
atcagtgatgatcggcgtttttgtttgcgtgggatttatgccggacgaaaaaaacgccatcgcgacgggctcat
tcggctatcagagcttcggcaactgcccgatacgccccatcatttcagtaacgatctgcaggtccagcaagaact
```

ggtcgacggtcttgaactctttattggtgtggccggtgtacttaacgtcgagcctggccaggccgaactgcacgc
cattgggcaggtcatgcacggaggttgagccggcagaggcgccgaattttcgcggcatgtcgaggttttcgctgg
cgacgtccagcaacgccttgacccactcaccctcagggtcgcggagcatcggttcggtcatcaagtagtccacgt
ccattgcaatggcgttcttcgtgctccaggcgcccaacttgtcggcaatctcggctttcagcacgtccaagcgtt
ttccctttggtacgcgtaggttgaccgtcagcgtgaaggtcttttcatccatgccgataaaggtcagtgatgtag
tcagaggccccatgaatgcatccgaaaagccaatacccagtttgcctcccaggtaatccagcccccagttgtcgg
cagcgtagcgcgcggcatcggaaatagggttttgcttgagtgcgatgttcccgtccaggctgtggatgaagccca
acatcctggcgacggggttcacaccggattcgggttctgatgaatgcgcagacacgccggtaacggtcagttgga
cttccttgc…

>delEGEG (420 bp removed: bases 2028652-2029071, inclusive)
…gtgacctgaccatgtagggcttcgagcaggaagcgcacgtccttggccttggcttcggtctggatcggcgcggg
tgggttgttcaccgggatcgattcgccctgagccaccagcgtattgagttcggcgatgcgggccttgatggggc
gctggtcttgttgaggttcaacaggcgctggttgacgttgagttccagtcgggtaagcaacaacttgagcgccgg
ggacatcaactggccggggaaggtctcggtgatttcaccgcaacggcgcaggcgatcgttaagctcgatctcggt
gcgcttcttttccagcttgttgttttccaccacgtggttcatatagccaatggcgatcagtattacgatcccggc
tattaccagcagggtgatcatgagtggtgtcaccggtgtgacctctttatagggtttacagtggagtgtagtgac
tgggccatccggcggatagggctgatcgaccagttgcccaggtagtttgcttctctatagataggcgtccgttac
atggatgcacattgccatcatttgtcggagcgaactatagcgccttgtcggacgccagaatataggcgccaaagc
ctgggcgggcacaatcctcccataagcccggaaagcagggcgaagtcattgatttaaataaatttatccttgggg
gttgacgacctttcaatccatccatagaatgcgcgccacttacagcgtaaagcacacagcgaaacgcggtaggga
gtgaatgttgtacgtgcaacgaaaacgccgatcagtgatgatcggcgttttgtttgcgtgggatttatgccgga
cgaaaaaaaacgccatcgcgacgggctcattcggctatcagagcttcggcaactgcccgatacgccccatcattt
cagtaacgatctgcaggtccagcaagaactggtcgacggtcttgaactctttattggtgtggccggtgtacttaa
cgtcgagcctggccaggccgaactgcacgccattgggcaggtcatgcacggaggttgagccggcagaggcgccga
attttcgcggcatgtcgaggttttcgctggcgacgtccagcaacgccttgacccactcaccctcagggtcgcgga
gcatcggttcggtcatcaagtagtccacgtccattgcaatggcgttcttcgtgctccaggcgcccaacttgtcgg
caatctcggctttcagcacgtccaagcgttttccctttggtacgcgtaggttgaccgtcagcgtgaaggtctttt
catccatgccgataaaggtcagtgatgtagtcagaggccccatgaatgcatccgaaaagccaatacccagtttgc
ctcccaggtaatccagcccccagttgtcggcagcgtagcgcgcggcatcggaaatagggttttgcttgagtgcga
tgttcccgtccaggctgtggatgaagcccaacatcctggcgacggggttcacaccggattcgggttctgatgaat
gcgcagacacgccggtaacggtcagttggacttccttgc…

*gluTTC* gene copies
*glyGCC* gene copies
IR82 inverted repeat
IR83 two identical copies of inverted repeat
IR43 inverted repeat
IR44 inverted repeat
delEGEG_1 primer binding site
delEGEG_2 reverse complement of primer binding site
delEGEG_3 primer binding site
delEGEG_4 reverse complement of primer binding site
delEGEG_f primer binding site
delEGEG_r reverse complement of primer binding site
delEGEG_f2 primer binding site
delEGEG_r2 reverse complement of primer binding site

#### *1.2 Re-insertion of* EGEG *tRNA locus into ΔEGEG (construction of rWT)*

To construct the 'reconstructed wildtype' (rWT) strain, the 420-bp encompassing the four-gene tRNA locus *gluTTC-glyGCC-gluTTC-glyGCC* (*EGEG*) was re-inserted into ΔEGEG at its wildtype chromosomal position. The aim is to construct a genotype that is isogenic to the original wildtype. This genotype was constructed to test whether any observed phenotypic effects are due to deletion of the four tRNA genes, rather than (i) any secondary genomic mutation(s) that may have been acquired during the construction process, or (ii) any lasting phenotypic effects of the engineering process.

Outline of the construction process:

1. Primers delEGEG_1 and delEGEG_4 were used to amplify a 1,603-bp fragment from SBW25 wildtype. The fragment was ligated into the pCR™8/GW/TOPO® vector and TA Cloning® kit

(Invitrogen™), and the sequence checked by Sanger sequencing with commercial primers GW1 and/or GW2.

2. Following the same protocol as in section 1.1 above, the confirmed fragment was transferred from pCR™8/GW/TOPO® into pUIC3, and exchanged with the homologous chromosomal fragment in ΔEGEG using scar-free, two-step allelic exchange.
3. The sequence of the manipulated locus in the resulting genotype, rWT, was confirmed by PCR amplification and Sanger sequencing with delEGEG_f/r.

```
>wildtype (shown here: bases 2027887-2029844)
…gtgacctgaccatgtagggcttcgagcaggaagcgcacgtccttggccttggcttcggtctggatcggcgcggg
tgggttgttcaccgggatcgattcgccctgagccaccagcgtattgagttcggcgatgcgggccttgatggggc
gctggtcttgttgaggttcaacaggcgctggttgacgttgagttccagtcgggtaagcaacaacttgagcgccgg
ggacatcaactggccggggaaggtctcggtgatttcaccgcaacggcgcaggcgatcgttaagctcgatctcggt
gcgcttcttttccagcttgttgttttccaccacgtggttcatatagccaatggcgatcagtattacgatcccggc
tattaccagcagggtgatcatgagtggtgtcaccggtgtgacctctttatagggtttacagtggagtgtagtgac
tgggccatccggcggatagggctgatcgaccagttgcccaggtagtttgcttctctatagataggcgtccgttac
atggatgcacattgccatcatttgtcggagcgaactatagcgccttgtcggacgccagaatataggcgccaaagc
ctgggcgggcacaatcctcccataagcccggaaagcagggcgaagtcattgatttaaataaatttatccttgggg
gttgacgacctttcaatccatccatagaatgcgcgccacttacagcgtaaagcacacagcgaaacgcggtaggga
gtgaatgttgtacgtgtgtccccttcgtctagtggcctaggacaccgccctttcacggcggtaacaggggttcga
gtcccctaggggacgccaatatgcgggaatagctcagttggtagagcacgaccttgccaaggtcggggtcgcgag
ttcgagtctcgtttcccgctccaaatttaagcggtattgccttcgggtggtactgagtgaaaccagggcataatc
ttcggatttaatctctggacactgaaatacacaccatgtgtttcagttgtgtgtccccttcgtctagtggcctag
gacaccgccctttcacggcggtaacaggggttcgagtcccctaggggacgccatttgcgggaatagctcagttgg
tagagcacgaccttgccaaggtcggggtcgcgagttcgagtctcgtttcccgctccatattcaacgaaaacgccg
atcagtgatgatcggcgtttttgtttgcgtgggatttatgccggacgaaaaaaaacgccatcgcgacgggctcat
tcggctatcagagcttcggcaactgcccgatacgccccatcatttcagtaacgatctgcaggtccagcaagaact
ggtcgacggtcttgaactctttattggtgtggccggtgtacttaacgtcgagcctggccaggccgaactgcacgc
cattgggcaggtcatgcacggaggttgagccggcagaggcgccgaattttcgcggcatgtcgaggttttcgctgg
cgacgtccagcaacgccttgacccactcaccctcagggtcgcggagcatcggttcggtcatcaagtagtccacgt
ccattgcaatggcgttcttcgtgctccaggcgcccaacttgtcggcaatctcggctttcagcacgtccaagcgtt
ttccctttggtacgcgtaggttgaccgtcagcgtgaaggtcttttcatccatgccgataaaggtcagtgatgtag
tcagaggccccatgaatgcatccgaaaagccaatacccagtttgcctcccaggtaatccagcccccagttgtcgg
cagcgtagcgcgcggcatcggaaatagggttttgcttgagtgcgatgttcccgtccaggctgtggatgaagccca
acatcctggcgacggggttcacaccggattcgggttctgatgaatgcgcagacacgccggtaacggtcagttgga
cttccttgc…
```

*gluTTC* gene copies
*glyGCC* gene copies
IR82 inverted repeat
IR83 two identical copies of inverted repeat
IR43 inverted repeat
IR44 inverted repeat
delEGEG_1 primer binding site
delEGEG_4 reverse complement of primer binding site
delEGEG_f primer binding site
delEGEG_r reverse complement of primer binding site

##### 1.3 *Deleting the lone copy of* glyGCC *from SBW25 (construction of Δ*glyGCC*)*

In engineered mutant Δ*glyGCC*, 82-bp encompassing the single-gene *glyGCC* locus at ~2,375,800 are removed from the wildtype genome. As before, the construction process used is scar-free (*i.e.*, no other trace of the construction process, such as antibiotic markers, are left in the genome of the deletion mutant).

Outline of the deletion process:

1. Primers delglyGCC_1, 2, 3, and 4 were used to construct a 965-bp deletion fragment via SOE-PCR (Ho et al. 1989). The fragment was ligated into the pCR™8/GW/TOPO® vector and TA

Cloning® kit (Invitrogen™), and the sequence checked by Sanger sequencing with commercial primers GW1 and/or GW2.

2. The confirmed deletion fragment was transferred from pCR™8/GW/TOPO® into the pUIC3 vector (Rainey 1999), using the *Bgl*II restriction sites incorporated into the deletion fragment construction primers.
3. The deletion fragment in the pUIC3 vector was exchanged with the homologous chromosomal fragment using scar-free, two-step allelic exchange. The sequence of the manipulated locus in the resulting genotype, Δ*glyGCC*, was confirmed by PCR amplification and Sanger sequencing with delglyGCC_f/r.

| Primer Name | Sequence (5'→3')[a] | Use |
|---|---|---|
| delglyGCC_1 | gaagatctTGGCCGACAAACTCATGGTG | Deletion fragment construction |
| delglyGCC_2 | GTCTTGGTGATGCGCATTCT | Deletion fragment construction |
| delglyGCC_3 | gaatgcgcatcaccaagacGATCCACAGATGTCCACTGGAC | Deletion fragment construction |
| delglyGCC_4 | gaagatctTCGTTGACCCAAGTGCCGATC | Deletion fragment construction |
| delglyGCC_f | ATGGCTTCCAGTTCGGTGTTC | Final PCR & sequencing |
| delglyGCC_r | TTGTCGGCTTATTGGCGGAG | Final PCR & sequencing |

[a]underline denotes *Bgl*II restriction sites (these are used for the cloning process, and are not included in the final, scar-free deletion). Capital letters denote the region annealing to the SBW25 genome sequence, and small letters denote non-annealing primer sequence. Highlighting relates to the nucleotide sequences below. Primers are also listed in Supplementary Table S1.

```
>wildtype (bases shown here: 2375271-2376519)
…atggcttccagttcggtgttcgcgtttgcggccgccaccgactggctcgacggctacctggcccgccgcctgga
acagagcacgcccttcggcgcgttcctcgacccggtggccgacaaactcatggtggccgttgcgctggtcctgtt
ggtgcaggaacacggcaacctgtggctgaccctgccggccgccgtgatcatcggccgtgagatagtggtgtcggc
cctgcgcgaatggatggccgaaatcggcgcccgcgcccacgtcgccgtctccaacatgggcaaatggaaaaccgc
cgcgcagatgctcgcgctggtcatcctgctggccaacccgtcggacttcaccttctgggtcctgctgggctacgc
cttgctgctcatcgccgccggcctgaccttgtggtccatgctccagtatctgcgcgccgcctggccgcatctgcg
taccaccgttgaaaagaaataaaactttttgaatcaaggggttgacgggttctgataattctatagaatgcgca
tcaccaagacgcgggaatagctcagttggtagagcacgaccttgccaaggtcggggtcgcgagttcgagtctcgt
ttcccgctccaaaatttgatccacagatgtccactggacgctggggtccacgaaatagacgcttaggcgtctttt
ttcgtttctgcggaaaaccactgggaaccagggtaatccggtacgaacaaggacaggattaagtacaacagacgg
gcccgttcgagtccgtattgtaactcgctccctcgggcctccagacgagcatcgggtgttggctctccaacagga
gctcgccatgcctctcttagaattcaccactcggcatgctcgaactattggtaaaaactacaccctcggcgactt
cgacggtctatccctctgggtgaccaaagcgggcggtaagctgtggctgttccgctactattgggaagggcgcca
gcaacgcatgtcctttggggcgtatcctgacgttagcctgaaacacgcacgtgagcgtcgtgacgaagcgcgcga
gctattggccgaaaatatcaacccttgcgcgcaccgaaagcaggcgcgccaagaagctgaagccgaagtaaagtg
ggcaattttagatcggcacttgggtcaacgatttactgatcattgcgatccgacgatccaagcggtttgcttac
caaaaaatcagctaagtcctcagcagtttcctccgccaataagccgacaa…
```

```
>delglyGCC (82-bp removed - bases 2375805-237588, inclusive)
…atggcttccagttcggtgttcgcgtttgcggccgccaccgactggctcgacggctacctggcccgccgcctgga
acagagcacgcccttcggcgcgttcctcgacccggtggccgacaaactcatggtggccgttgcgctggtcctgtt
ggtgcaggaacacggcaacctgtggctgaccctgccggccgccgtgatcatcggccgtgagatagtggtgtcggc
cctgcgcgaatggatggccgaaatcggcgcccgcgcccacgtcgccgtctccaacatgggcaaatggaaaaccgc
cgcgcagatgctcgcgctggtcatcctgctggccaacccgtcggacttcaccttctgggtcctgctgggctacgc
cttgctgctcatcgccgccggcctgaccttgtggtccatgctccagtatctgcgcgccgcctggccgcatctgcg
taccaccgttgaaaagaaataaaactttttgaatcaaggggttgacgggttctgataattctatagaatgcgca
tcaccaagacgatccacagatgtccactggacgctggggtccacgaaatagacgcttaggcgtcttttttcgttt
ctgcggaaaaccactgggaaccagggtaatccggtacgaacaaggacaggattaagtacaacagacgggcccgtt
cgagtccgtattgtaactcgctccctcgggcctccagacgagcatcgggtgttggctctccaacaggagctcgcc
atgcctctcttagaattcaccactcggcatgctcgaactattggtaaaaactacaccctcggcgacttcgacggt
ctatccctctgggtgaccaaagcgggcggtaagctgtggctgttccgctactattgggaagggcgccagcaacgc
atgtcctttggggcgtatcctgacgttagcctgaaacacgcacgtgagcgtcgtgacgaagcgcgcgagctattg
gccgaaaatatcaacccttgcgcgcaccgaaagcaggcgcgccaagaagctgaagccgaagtaaagtgggcaat
tttagatcggcacttgggtcaacgatttactgatcattgcgatccgacgatccaagcggtttgcttaccaaaaaa
tcagctaagtcctcagcagtttcctccgccaataagccgacaa…
```

taa stop codon of *pgsA* (PGP synthase)

atg start codon of *pflu2192* (pseudogene)
*glyGCC* single copy tRNA gene encoding tRNA-Gly(GCC)
delG_1 primer binding site
delG_2 reverse complement of primer binding site
delG_3 primer binding site
delG_4 reverse complement of primer binding site
delG_f primer binding site
delG_r reverse complement of primer binding site

##### *1.4 Isolation of ΔEGEG-**rplC** from M5-1*

Genotype M5-1 carries two mutations: a large-scale (~1 Mb) duplication, and a non-synonymous mutation in *rplC*, the gene encoding the 50S ribosomal protein L3 (c128t, causing amino acid change T43I). We would like to assess the effect of each of these mutations alone.

- Any of M1-1, M2-1, M4-1 can be used to assess the effect of the ~1 Mb duplication fragment alone; each carries a similar duplication fragment to M5-1, without any other mutations (see Supplementary Table S2).
- To obtain the *rplC* mutation alone we leveraged the instability of the large duplication fragment in M5-1. M5-1 was grown up in liquid culture (28˚C, 200 rpm, ~16 hours, KB), and dilution plated on KB agar (28˚C, ~30 hours). The resulting colonies were a mixture of large and small types. Five large and five small colonies were purified, grown up in liquid KB (28˚C, 200 rpm), and stored at -80˚C. Analyses revealed (i) amplification of the M5-1 duplication fragment junction in all large colony-forming strains, but none of the five small colony-forming strains (PCR with primers M5_junct_f/r; see Supplementary Table S1), and (ii) retention of the *rplC* mutation was confirmed in eight of the ten strains (by PCR and Sanger sequencing with primers rpsJ_f/rplC_r; see below). In the final two strains (M5-1-Large-3 and M5-1-Small-3), the *rplC* Sanger sequencing reactions were unsuccessful and hence the *rplC* allele was not identified in either. One small colony-forming strain, M5-1-Small-4 (hereafter called ΔEGEG-*rplC*), was used for further work.
- Together, these results are consistent with the small colony-forming types carrying only the *rplC* mutation (and not the duplication fragment).

```
>wildtype rplC region (6047892 - 6048724; reverse strand)
…gatccgtactcataagcgcgtactggacatcgtccagccaacggataaaaccgttgatgcacttatgaagctcg
atctggcggccggtgtggaagtacagatcagcctcggctaagacttgggtcttagtcgtgtaacgctctgaaatg
ggcggccatagcgggtgaaagccccgtacactcatgaggtttacaacatgactattggtgtagtcggtcgtaaat
gcggtatgacccgtattttcaccgaagaaggtgtctccattccggtcacggtcattgagatcgaaccgaatcgcg
tcacccagttcaaaactgaagagaccgatggctatcgtgcagtgcaagtcactgtcggcgagcgtcgtgcttcgc
gcgtgactgctgctcaagcaggtcacttcgctaaagcaaacgttgcagctggtcgcactgttatggagttccgtc
tcgaagacggcgactaccaggctggcgatctgatcaacgctgaaatcttcgccgctggtcaactggttgatgtaa
ccggtcagtccaaaggtaaaggcttccagggtacgatcaagcgttggaatttccgtggccaagacaacactcacg
gtaactccgtttcccaccgcgtcccgggctctattggccagtgccagactcctggtcgtgtattcaagggcaaaa
aaatgtccggtcatatgggcgctgagcgcgtgaccgtgcagtccctcgaagtagtgcgcgtcgacgctgaacgca
atctgttgttggtcaagggtgctgttcctggcgctactggcggcaacctggttgtacgtccagcggccaaggctc
gcggttaag…
```

*rplC* (636 bp, 211 amino acids)
c128t (*rplC* mutation in M5-1 and SBW25-*rplC*)
GATCCGTACTCATAAGCGCGTACTG rpsJ_f; primer binding site
CCTTGAATACACGACCAGGAGTCTG rplC_r; reverse complement of primer binding site

**Supplementary Text S2**

**Identification of emergent junctions and duplication fragments in genotypes isolated from day-21 of the evolution experiment**

Five genotypes were isolated from day-21 of the evolution experiment: one from each mutant line. These genotypes are M1-1, M2-1, M3-1, M4-1, M5-1. Each was predicted by whole genome re-sequencing to carry a large-scale duplication fragment of up to ~1 Mb in size (see main manuscript Table 1, Figure 4). In this supplementary text, we outline the computational and laboratory methods used to predict and confirm the molecular details of each of the five duplication fragments.

#### *2.1 Methods of duplication fragment/junction identification in isolates*

The segments of duplicated genomic DNA in genotypes M1-1 to M5-1 (see Table 1, main manuscript) were identified using computational and lab-based techniques. The broad strategy for identification of the duplication mutation in each isolate was:

*Bioinformatic analysis led to proposed duplication fragments in each isolate:*

1. Standard analysis of the NGS data with *breseq* and Geneious (v11.1.4).
   ⇒ Identification of predicted duplication fragments in M1-1 and M3-1
2. *De novo* alignment of unused reads from the Geneious alignment, searching specifically for unique contigs covering the duplication junction.
   ⇒ Further evidence for proposed duplication fragments in M1-1 and M3-1
3. Close analysis of the coverage data from the Geneious alignment.
   ⇒ Approximate duplication region identified in all five genotypes
4. Manual inspection of both ends of the duplication region in Geneious alignments.
   ⇒ Identification of a proposed duplication fragment in M2-1, M4-1, and M5-1
5. Demonstration of significant numbers of raw reads aligning to the unique duplication junction proposed for each genotype (and 0 for the same alignment performed with the raw reads of other genotypes) (see Tables 2.1 and 2.2 below).

*Laboratory-based confirmation of the proposed duplication fragments in each genotype:*

1. PCR primers were designed and used to amplify the unique duplication junction proposed for each genotype (alongside negative controls; see below for details of PCR fragments).
   ⇒ Expected PCR products were obtained in M1-1, M3-1, M4-1, M5-1 (see Fig. 2.1 below).
2. The PCR products above were cloned into a sequencing vector (pCR™8/GW/TOPO™, Invitrogen™), and Sanger sequenced to confirm the sequence of the duplication junctions.
   ⇒ The exact junction was confirmed in M1-1, M3-1, and M5-1
   ⇒ The junctions in M2-1 and M4-1 were too repetitive (due to within-gene *rhs* repeats) to sequence fully, each side of the cloned fragment was shown to belong uniquely to one side of the proposed junction.

#### *2.2 Duplication junction sequences*

This section lists the unique, emergent duplication junction sequence proposed for each of the five evolution experiment isolates (M1-1 to M5-1). These are the sequences to which the raw reads from the relevant genotype were aligned to obtain the read numbers listed in Table 2.1 below (footnote *b*).

```
>M1-1_30bp_junct
TCACCTTTGCGGGTGGTGAGACCGCTATGC

>M2-1_46bp_junct
GTTGccatcaggcagggtttttcttacgagccgaccggcGTGGTCT
```

```
>M3-1_88bp_junct
GCCGTCCACCTATCgTGGGAGGGGGCTTGCCCCCGATAGCGGTGTGTCAGTCGATTCATCAGGTGACTGATTCAG
CGCCATCGGGGCG

>M4-1_714bp_junct (shared region = 696 bp)
AACAAACGcacgttgtcgtagcggtagttgacctggctgccatcggcgttgaggcgacggctgatcaggtgcaag
ccgtcggcgtattcgtagcgggtaacattccccagctcatcgcgttcggaagtgatttttccgtaggggttgtag
ctgtattccctgagcgcaccgcccgatagcacaacgccaatcaaccgcccaacactgtcccattgatactgggtc
agcgccccatgctcatcctcacgcgccacctgccggccaagatcgtcatagcgataacgcttgatcccaccattc
ggcaactgctcttcaagcaactgcccacgttcattccaaaccaacttctgacagctgtgatccggataccaaacc
ccaaccagttgcccatatttgttgtagctgtagtcagtagcgtgaccatcaggatcggtcctacgcgtcacatcg
ccctgatcattgcgctcatacttccaaaccgcttcaccacgccttacgacccgtacgaacccattgtcatgctcg
taggtcgtcggctcatcctccccggaaacaacgccaccaagcgtccagcgtcgtcgtactgatacgccgtcacc
gccccagcgggtcctgcttaaccgtcagccggcctttttcatcgtaagatttaaagtgctcggcaccgtccgga
tccacccgctgcaccagccgcgcccgctgGTCATGCACG

>M5-1_75bp_junct:
AATCGCCGCTTCCgccaaactattggtaatcgcaggccgagaaaccgtaggcaagggcagggCAGTCGACCGATT
```

Duplication junctions key:
SIDE 1 of each junction
SIDE 2 of each junction
bases occurring on both side 1 and side 2 of the junction (*i.e.*, repeats)
*Note 1: each side (+ grey lettered shared region, where present) is a unique sequence in the wildtype genome sequence.
**Note 2: no inversions; both sides of each junction occur on the same strand, in the same direction.
***Note 3: if the sides are highly similar, underline indicates bases that differ between side.

| Geno-type | Size | | | Computational evidence | | | Laboratory evidence[c] |
|---|---|---|---|---|---|---|---|
| | bp | #duplicated genes[a] | #duplicated tRNA genes[a] | Evidence type | Fold coverage | Reads[b] | |
| M1-1 | 1,016,166 | 916 | 7 | Called by breseq, coverage plots, bin/fragment alignments | 1.5 | 13 | PCR |
| M2-1 | 1,012,338 | 910 | 7 | Coverage plots, bin/fragment alignments | 1.4 | 24 | PCR |
| M3-1 | 489,709 | 425 | 1 | Called by breseq, coverage plots, bin/fragment alignments | 1.8 | 21 | PCR, Sanger |
| M4-1 | ~1,012,000 | 910 | 7 | Coverage plots, fragment alignments | 1.3 | n/a | PCR |
| M5-1 | 1,012,339 | 910 | 7 | Coverage plots, fragment alignments | 1.6 | 16 | PCR, Sanger |

**Table 2.1. Features of the large-scale duplication fragments.** [a]Lists of duplicated genes, including tRNA genes, can be found in Supplementary Table S3. [b]The number of raw reads found spanning the entire, unique duplication junction fragment in each genotype. Duplication junction sequences are listed below. Note that in order to reach unique sequence, the duplication junction in M4-1 extends >700 bp, and thus cannot be covered by single raw reads (which are a maximum length of 250 bp). [c]Lab evidence is in more detail in section 2.3.

| Junction name | Junction seq. length (bp) | Number of raw reads covering entire junction sequence with no mismatches | | | | | |
|---|---|---|---|---|---|---|---|
| | | M1-1 | M2-1 | M3-1 | M4-1 | M5-1 | W1-1 |
| M1-1_30bp_junct | 30 | **21** | 0 | 0 | 0 | 0 | 0 |
| M2-1_46bp_junct | 46 | 0 | **77** | 0 | 0 | 0 | 0 |
| M3-1_88bp_junct | 88 | 0 | 0 | **21** | 0 | 0 | 0 |
| M4-1_714bp_junct | 714 | Junction sequence too long for unbroken coverage by 250 bp reads | | | | | |
| M5-1_75bp_junct | 75 | 1 | 0 | 0 | 0 | **16** | 0 |

**Table 2.2. Coverage of the emergent duplication junction identified in each isolate.** Each unique, emergent duplication junction sequence (first column) was used as a reference to which the raw reads obtained during

genome re-sequencing of six isolates (M1-1, M2-1, M3-1, M4-1, M5-1, W1-1) were aligned. Numbers of raw reads aligning perfectly (no mismatches and unbroken coverage) to each reference sequence, for each isolate, is listed. **Bold**=isolate in which the junction was identified; matches are notably absent in the other isolates (with the exception of 1 read from M1-1 matching M5-1_75bp_junct). This result provides evidence that each of the five duplication junctions (and therefore, duplication fragments) is distinct from the others.

##### *2.3 Duplication fragments amplified by PCR, ligated into pCR$^{TM}$8/GW/TOPO$^{TM}$*

This section lists the PCR products, including primer sequences, amplified to confirm each of the five emergent duplication junctions in section 2.2 above. Details of the PCR conditions used in each case are provided in Table 2.3 (below), and Figure 2.1 (below) shows the PCR products obtained.

```
>M1-1_emergentjunction_PCR (658 bp; base 2,864810 sewn to 1,848,645)
GAGCCAGTGGACGAAAGCTGTCCGGTTCATAGAGGTAGCTGCGGTGACGGTCCGCATGGTGTTCGGCGATGAGTG
TGTCGCCTTGCCAGAAGAACTCGGTGGTTACGTCGTCGACGGTTTTGCTGATACGCCGCCCAAACGGGTCGTAGC
GGTAGCTGGCGGTCTGGCCGTTAGGTTGGGTGATGCCGATCAGCCGATGCTGGCAGTCGTAGCGGTATTCGGTGA
CAAGTGCGTGGCCTTTGCCGCGGCGTTCGCGGATGAGGTTGCCGAAGGCGTCGTAGTCGTAGTGGTGGTCGCCTT
GGATCATCAGGCGGTTGCCGGCGACGATGTCGGGGCCGGGACGGTCTTGCATGAGCAGGTTGCCGGCGGGGTCGT
GGCCGAAGCGTTCTGGCTCGTCTTGGGAGTGGTTGGCGCGGGTGAGGCGGCTTAGTGGGTCATAGTGGTAGTGCT
GTTCACCTTTGCGGGTGGTGAGACCGCTATGCGCTTTAGTGATTTGCTCGATGCCGCCCGTAGCAACCCGCTGGA
TGTCACCATCCCCGCCGAATGGGCCCAGGGTCGCGCGACCTTTGGTGGGTTGGTCGCCGCATTGCAATACGAAGC
CCTGCGTGCCCAAGTGCCGGCCGATCGCCCGCTGCGGTCGCTGGCAATCACCTTCGTC

>M2-1_emergentjunction_PCR (2958 bp; base 2,865,277 sewn to 1,852,939)
GAGCCAGTGGACGAAAGCTGTCCGGTTCATAGAGGTAGCTGCGGTGACGGTCCGCATGGTGTTCGGCGATGAGTG
TGTCGCCTTGCCAGAAGAACTCGGTGGTTACGTCGTCGACGGTTTTGCTGATACGCCGCCCAAACGGGTCGTAGC
GGTAGCTGGCGGTCTGGCCGTTAGGTTGGGTGATGCCGATCAGCCGATGCTGGCAGTCGTAGCGGTATTCGGTGA
CAAGTGCGTGGCCTTTGCCGCGGCGTTCGCGGATGAGGTTGCCGAAGGCGTCGTAGTCGTAGTGGTGGTCGCCTT
GGATCATCAGGCGGTTGCCGGCGACGATGTCGGGGCCGGGACGGTCTTGCATGAGCAGGTTGCCGGCGGGGTCGT
GGCCGAAGCGTTCTGGCTCGTCTTGGGAGTGGTTGGCGCGGGTGAGGCGGCTTAGTGGGTCATAGTGGTAGTGCT
GTTCACCTTTGCGGGTGTCGAGCAGGCGGGTGAGGTTGCCGGATTTGTCGTAGTCGTAATGGCGTTGGTAGAGGG
TGTATTCCGGTTGGGTGACGGCGTGGGCGTGCAAGCGTTGCTGGTCGTCATAGTGGTAGTGGCTGATCAGTTGGC
CTTGTTGGCGTTGGTGTTCCTGGCCGGCTTTGTAGAGATGCGAAGTGAGGATCTCGCCGTTCAGTTCGACGGTGG
CCAGGTGGCCGCCTTTGTCGTGGTTGAAGGTGAGGCGGTTGTTGTCGGGCAGGCGCAGGTTTGTTAGTTGGCCGC
AGTTGTCGTAGCCGTAGCGCAGGGTGCCCCAGCCTTGGTGTTCGGCGGTGAGGCGGTTTTGGGCGTCGTATTCGT
AAGCCAGGGCCCAATGGCCATCTTCGACGCTGAGGAGGTTGCCCTGGCGGTCGTAGGCGTAATCAACGTTGTTGc
catcaggcagggtttttcttacgagccgaccggcGTGGTCTCGCTCGTAGCGGGTAACCAACTGACTGCCGTCAT
CGCCGTGTTCCGTCTTTTCCTGAAGGTTGCCGTTGAGGTCGTAAACGTAGGCCGTGCGCTGACCGTCAAACCCGG
TTTCCTGCTGGATCAGGCCATTGCTGTGGTACTGAAGTCGGTAGGTCTCGCCAACCTCGTTTTCGATTTCGGTCA
GCAACAAACCCACGTTGTCGTAGCGGTAGTTGACCTGGCTGCCATCGGCGTTGAGGCGACGGCTGATCAGGTGCA
AGCCGTCGGCGTATTCGTAGCGGGTAACATTCCCCAGCTCATCGCGTTCGGAAGTGATTTTTCCGTAGGGGTTGT
AGCTGTATTCCCTGAGCGCACCGCCCGATAGCACAACGCCAATCAACCGCCCAACACTGTCCCATTGATACTGGG
TCAGCGCCCCATGCTCATCCTCACGCGCCACCTGCCGGCCAAGATCGTCATAGCGATAACGCTTGATCCCACCAT
TCGGCAACTGCTCTTCAAGCAACTGCCCACGTTCATTCCAAACCAACTTCTGACAGCTGTGATCCGGATACCAAA
CCCCAACCAGTTGCCCATATTTGTTGTAGCTGTAGTCAGTAGCGTGACCATCAGGATCGGTCCTACGCGTCACAT
CGCCCTGATCATTGCGCTCATACTTCCAAACCGCTTCACCACGCCTTACGACCCGTACGAACCCATTGTCATGCT
CGTAGGTCGTCGGCTCATCCTCCCCCGGAAACAACGCCACCAAGCGTCCAGCGTCGTCGTACTGATACGCCGTCA
CCGCCCCCAGCGGGTCCTGCTTAACCGTCAGCCGGCCTTTTTCATCGTAAGATTTAAAGTGCTCGGCACCGTCCG
GATCCACCCGCTGCACCAGCCGCGCCCGCTGGTCATGCACGTAGACTTCCTGACTGCCGTCGGCGTTAAACACCG
TGACCTGCCCGTTGTCATCCCAGGCATACCGCGTGTCCATCTGCGAAAAACTGGCCCAGTGCCGGACACACCTCG
CCGCCTTGCCTGACCTTTCCCACTCCCAGTAAAAACTCGCCCCACCGGCCAAACCACGCTCAAGAATGACGTGCT
GTTCGTCGTACCGATAAACCTCGCGTTCACCGACAGCATTGGTCGCAGAAACCAGTCGCCCAGCACCGTCATACG
CGTAGGAAACAACGTTCTGTTCCGTCACCCAGACAAAAGGCTCAAGACCCGTGGCCCGATGGATCTGATAATCCA
CCGCCACGATCCGCCCCGACGCATAACGCAAAAACAACGAACGCCCGACGCCGTTATCCAGCCGCTCAATCTGCC
CCAGATAATTGCGGGAAATGCGCAACCGGTTGTCATACGCATCGCTGATCGCCGTCAGTACACCGTCGCGAAAGT
GGTAGAACCGTGACGACTGAGCCAGCACCAGTTCATCAGGCAACGACCCCAAGTAGATCGCCGCTTCAGCCAAAC
TATTGGTAATCGCAGGCCGAGAAACCGTAGGCAAGGGCAGGGCAGTCGACCGATTTTCATGATCTGTCCACACCA
CCGAATCACCGGAAACACTCAACCGATGCGCCAGCGAGTGACTCCAACCAAACCCCAGCCCACAATCCATTTCCA
CCGCACTGGTGCGGTACAAGCGCGTCCACTCAAACGGCAAAACCCCGTCCAACGCCCCATCGGCCAGCGTCAGCA
GTTCCTCGCCGGTGACCATCGACACCGGGCAACCGTTAGTGGCTGTTTTGTCCGAAGACGCGGCCGCCTCTCCAG
CCGGGTTCTTAGCAACAGCCGGCACGTCATCCACAGGCTCCTGCCGCACCAACACCGTCCGCTGCTTGGCCTTGT
CCCGAACCACCGGCACCGGGTTGGCAACCACCTGATCCCCAGCCTTCAACGGCACATCCGGAATCGTCTTGATCG
```

GCGTCCCCGCGCTGCCCAGCAATAACGGCTTGGCCGTTTCGACATGCGCTTCCAGCCGAGACCCGACCACTTGAT
CAGCCAGCCGCTCCAGCCATTTGCGTGCTCTAC

>M3-1_emergentjunction_PCR (986 bp; base 2,786,050 sewn to 2,96,499)
GTGTATGACGATGACGAGCAGAACCCGCGTGACCAGCAATTGGTCAACGACTACGGTTTTGCCTACACGCCGGGC
CGCCAGTGGCTGTACCCGGCCGACCTGCGGGTGCAGGGGCGCCTGGTCAATGACTTGCTGTTGAGCGTGCCGTCC
ACCTATCgTGGGAGGGGGCTTGCCCCCGATAGCGGTGTGTCAGTCGATTCATCAGGTGACTGATTCAGCGCCATC
GGGGCGTCGGTATCTACACAATTCTGGCTATGGCGCTGGACCTGTGGCGAGGGAGCTTGCTCCCGCTGACGGCCT
GACAGCTGACCGCCATCTAAATGTTACCCCGCGATCCAAATGTGGGAGGGGGCTTGCCCTCGATAGCGGTGTGTC
AGTCGATTCATCAGGTGACTGATTCAGCGCCATCGAGGCATCGGTATCTACACCACTCTGGCTATGGCGCTGGAC
CTGTGGCGAGGGAGCTTGCTCCCGTGGCGGCCTGACAGCTGACCTGGATGCTGGATCAGACCGGGTACATATCCG
TTATTTAGGTAACGGCCGCTATGGGTTCCGCTTTTACAGCGGCTCACTTTTGAAAAGCGCAAAAGTAAGCAAAAC
GCTCTTGCCCCAACACTCGGCACCTCGCTCACGCTCGGTGTGCCCGTAATCCGACAGGGATTTGGGGGGCCGCCG
CCACGCGCCATCCATGGCGCGGGGCGGCTAAACCGGCATCCCTGCCGGTTTACCCCCCAAATCCCTGTCGAATTC
CGGCCAGCGTGTTTGACGGGGCGCCTGAGATCAAAAGCGAGATCAAGATCAAGATCAAGAGCGGCTCGCTTCGCA
TCGTGGTTAGTGGTTGAGGGGCTGCACAGTTGTGTCGATAACTTTGGTTATCGCTGGTAAGCCGCCGTCTACAAG
ACTTGTAGTCTCTCGCCCAATCCGCCGTTATCCGCCAATGCCAGAAACTCATCCCCTAGCCGCCGGCTTTCGCTC
ATCGCTGTCTG

>M4-1_emergentjunction_PCR (~2958 bp; base 2,866,174 sewn to 1,853,836)
gagccagtggacgaaagctgtccggttcatagaggtagctgcggtgacggtccgcatggtgttcggcgatgagtg
tgtcgccttgccagaagaactcggtggttacgtcgtcgacggttttgctgatacgccgcccaaacgggtcgtagc
ggtagctggcggtctggccgttaggttgggtgatgccgatcagccgatgctggcagtcgtagcggtattcggtga
caagtgcgtggcctttgccgcggcgttcgcggatgaggttgccgaaggcgtcgtagtcgtagtggtggtcgcctt
ggatcatcaggcggttgccggcgacgatgtcggggccgggacggtcttgcatgagcaggttgccggcggggtcgt
ggccgaagcgttctggctcgtcttgggagtggttggcgcgggtgaggcggcttagtgggtcatagtggtagtgct
gttcacctttgcgggtgtcgagcaggcgggtgaggttgccggatttgtcgtagtcgtaatggcgttggtagaggg
tgtattccggttgggtgacggcgtgggcgtgcaagcgttgctggtcgtcatagtggtagtggctgatcagttggc
cttgttggcgttggtgttcctggccggctttgtagagatgcgaagtgaggatctcgccgttcagttcgacggtgg
ccaggtggccgcctttgtcgtggttgaaggtgaggcggttgttgtcgggcaggcgcaggtttgttagttggccgc
agttgtcgtagccgtagcgcagggtgccccagccttggtgttcggcggtgaggcggttttgggcgtcgtattcgt
aagccagggcccaatggccatcttcgacgctgaggaggttgccctggcggtcgtaggcgtaatcaacgttgttgc
catcaggcagggtttttcttacgagccgaccggcatggtcgcgctcgtagcgggtaaccaactgactgccgtcat
cgccgtgttcggtcttttcctgaagattgccgttgaggtcgtaaacgtaagcggtgcgctgaccgtcaaagccga
tctcctgttggatcaggccgttggcgtggtaatcaagcttgtaggtttcgccaacctcgttttcgatctcggtca
gcAACAAACGcacgttgtcgtagcggtagttgacctggctgccatcggcgttgaggcgacggctgatcaggtgca
agccgtcggcgtattcgtagcgggtaacattccccagctcatcgcgttcggaagtgatttttccgtaggggttgt
agctgtattccctgagcgcaccgcccgatagcacaacgccaatcaaccgcccaacactgtcccattgatactggg
tcagcgccccatgctcatcctcacgcgccacctgccggccaagatcgtcatagcgataacgcttgatcccaccat
tcggcaactgctcttcaagcaactgcccacgttcattccaaaccaacttctgacagctgtgatccggataccaaa
ccccaaccagttgcccatatttgttgtagctgtagtcagtagcgtgaccatcaggatcggtcctacgcgtcacat
cgccctgatcattgcgctcatacttccaaaccgcttcaccacgccttacgacccgtacgaacccattgtcatgct
cgtaggtcgtcggctcatcctcccccggaaacaacgccaccaagcgtccagcgtcgtcgtactgatacgccgtca
ccgcccccagcgggtcctgcttaaccgtcagccggccttttcatcgtaagatttaaagtgctcggcaccgtccg
gatccacccgctgcaccagccgcgcccgctgGTCATGCACGTAGACTTCCTGACTGCCGTCGGCGTTAAACACCG
TGACCTGCCCGTTGTCATCCCAGGCATACCGCGTGTCCATCTGCGAAAAACTGGCCCAGTGCCGGACACACCTCG
CCGCCTTGCCTGACCTTTCCCACTCCCAGTAAAAACTCGCCCCACCGGCCAAACCACGCTCAAGAATGACGTGCT
GTTCGTCGTACCGATAAACCTCGCGTTCACCGACAGCATTGGTCGCAGAAACCAGTCGCCCAGCACCGTCATACG
CGTAGGAAACAACGTTCTGTTCCGTCACCCAGACAAAAGGCTCAAGACCCGTGGCCCGATGGATCTGATAATCCA
CCGCCACGATCCGCCCCGACGCATAACGCAAAAACAACGAACGCCCGACGCCGTTATCCAGCCGCTCAATCTGCC
CCAGATAATTGCGGGAAATGCGCAACCGGTTGTCATACGCATCGCTGATCGCCGTCAGTACACCGTCGCGAAAGT
GGTAGAACCGTGACGACTGAGCCAGCACCAGTTCATCAGGCAACGACCCCAAGTAGATCGCCGCTTCAGCCAAAC
TATTGGTAATCGCAGGCCGAGAAACCGTAGGCAAGGGCAGGGCAGTCGACCGATTTTCATGATCTGTCCACACCA
CCGAATCACCGGAAACACTCAACCGATGCGCCAGCGAGTGACTCCAACCAAACCCCAGCCCACAATCCATTTCCA
CCGCACTGGTGCGGTACAAGCGCGTCCACTCAAACGGCAAAACCCCGTCCAACGCCCCATCGGCCAGCGTCAGCA
GTTCCTCGCCGGTGACCATCGACACCGGGCAACCGTTAGTGGCTGTTTTGTCCGAAGACGCGGCCGCCTCTCCAG
CCGGGTTCTTAGCAACAGCCGGCACGTCATCCACAGGCTCCTGCCGCACCAACACCGTCCGCTGCTTGGCCTTGT
CCCGAACCACCGGCACCGGGTTGGCAACCACCTGATCCCCAGCCTTCAACGGCACATCCGGAATCGTCTTGATCG
GCGTCCCCGCGCTGCCCAGCAATAACGGCTTGGCCGTTTCGACATGCGCTTCCAGCCGAGACCCGACCACTTGAT
CAGCCAGCCGCTCCAGCCATTTGCGTGCTCTAC

>M5-1_emergentjunction_PCR (611 bp; base 2,866,785 sewn to 1,854,447)
CAACACCAACTCATCGGGACAGCACCCAGATAAATCGCCGCTTCCgccaaactattggtaatcgcaggccgaga
aaccgtaggcaagggcagggCAGTCGACCGATTTTCATGATCTGTCCACACCACCGAATCACCGGAAACACTCAA

```
CCGATGCGCCAGCGAGTGACTCCAACCAAACCCCAGCCCACAATCCATTTCCACCGCACTGGTGCGGTACAAGCG
CGTCCACTCAAACGGCAAAACCCCGTCCAACGCCCCATCGGCCAGCGTCAGCAGTTCCTCGCCGGTGACCATCGA
CACCGGGCAACCGTTAGTGGCTGTTTTGTCCGAAGACGCGGCCGCCTCTCCAGCCGGGTTCTTAGCAACAGCCGG
CACGTCATCCACAGGCTCCTGCCGCACCAACACCGTCCGCTGCTTGGCCTTGTCCCGAACCACCGGCACCGGGTT
GGCAACCACCTGATCCCCAGCCTTCAACGGCACATCCGGAATCGTCTTGATCGGCGTCCCCGCGCTGCCCAGCAA
TAACGGCTTGGCCGTTTCGACATGCGCTTCCAGCCGAGACCCGACCACTTGATCAGCCAGCCGCTCCAGCCATTT
GCGTGCTCTAC
```

Key for PCR products to confirm duplication junctions:

```
FORWARD PRIMER see Table S2.2 and Supplementary Table S1
REVERSE PRIMER see Table S2.2 and Supplementary Table S1
SIDE 1 of each junction ; SIDE 2 of each junction
bases occurring on both side 1 and side 2 of the junction (i.e., repeats)
```

| Genotype | Forward primer | Reverse primer | Annealing temp (˚C) | Extension time (min) | $MgCl_2$ volume (µl) | Expected product size (bp) |
|---|---|---|---|---|---|---|
| M1-1 | M1and2_junct_f | M1_junct_r | 58 | 1 | 2.5 | 658 |
| M2-1 | M1and2_junct_f | M5_junct_r | 59 | 3 | 3.0 | ~3,000 |
| M3-1 | M3_junct_f | M3_junct_r | 58 | 1 | 2.5 | 986 |
| M4-1 | M1and2_junct_f | M5_junct_r | 59 | 3 | 2.5 | 2,958 |
| M5-1 | M5_junct_f | M5_junct_r | 58 | 1 | 3.0 | 611 |

**Table 2.3. PCR conditions for duplication junction amplification.** PCRs were performed using GoTaq® DNA Polymerase (Promega), according to the manufacturer's recipe (for a total volume of 25 µl) and conditions. All primer sequences are listed in Supplementary Table S1.

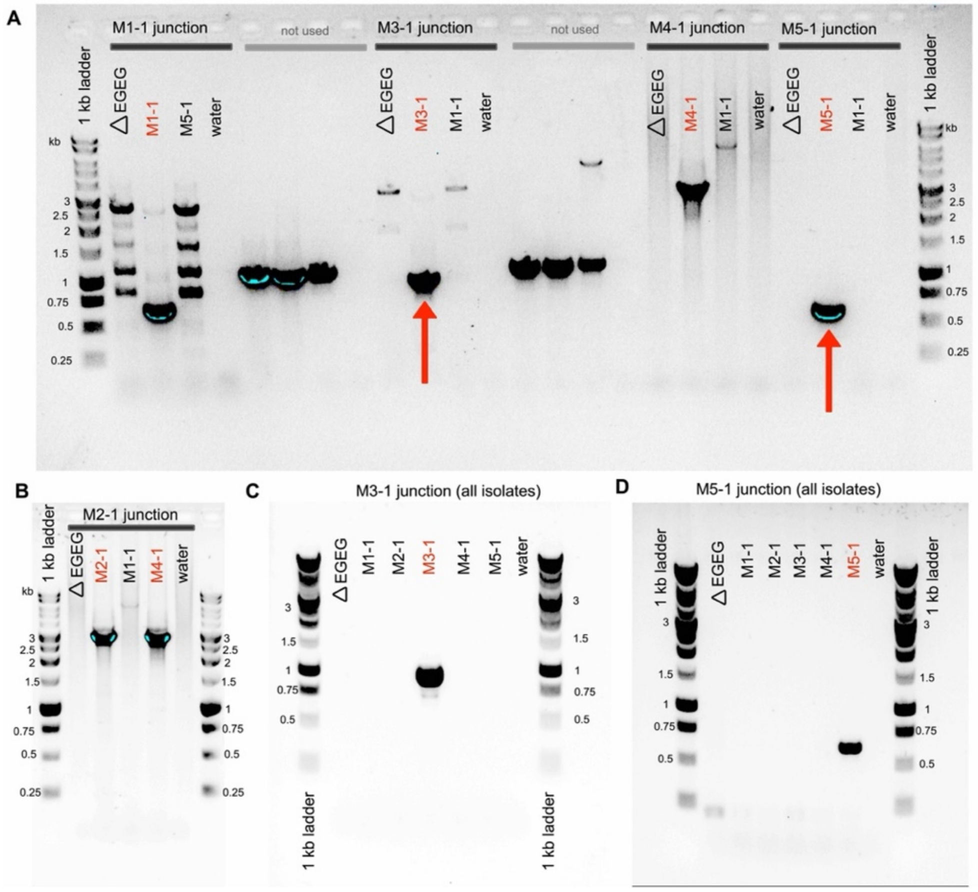


**Figure 2.1. Agarose gel images showing PCR products from duplication junction PCRs.** Red arrows indicate the emergent PCR bands for (**A**) M1-1, M3-1, M4-1, and M5-1 and (**B**) M2-1 and M4-1, versus absence of the same band in control reactions. Expected (and confirmed) band sizes for each evolved genotype are provided in Table 2.2 above. Gel photograph colours were inverted using Preview (11.0) to better detect faint PCR products. PCR products indicated by red arrows were purified, ligated into a sequencing vector (pCR$^{TM}$8/GW/TOPO$^{TM}$; Invitrogen$^{TM}$) and Sanger sequenced. (**C-D**) Duplication junction PCRs for M3f (panel C) and M5f (panel D) on all five mutant line isolates and controls, with both sets of PCRs showing specific duplication junction amplification. PCR conditions as listed in Table 2.2 above.

**Supplementary Text S3**
**Identification of emergent junctions and duplication fragments in day 28 populations**

Nine samples were used for population-level whole genome re-sequencing: day 28 populations from lines M1, M2, M3, W1, W2, W3, W4, W5, and a control SBW25 isolate. We note that the day 28 populations from M4 and M5 were excluded due to the discovery of a significant level of external contaminants (see Supplementary Figure S1). Indeed, a major motivation for sequencing the remaining day-28 populations was to rule out further contamination issues.

In this supplementary text, we outline the computational methods used to predict putative duplication fragments in each sequenced population.

#### 3.1 Methods of duplication fragment/junction identification in populations

The segments of duplicated genomic DNA in day 28 populations from lines M1, M2 and M3 (see main manuscript Table 2) were predicted using computational techniques. The broad strategy for identification of the duplication mutation in each population was:

1. Analysis of the NGS data with *breseq* (on polymorphism detection settings)
   ⇒ Identification of 9 predicted duplications fragments: M1d28-F1, M1d28-F3, M1d28-F4, M2d28-F2, M2d28-F3, M2d28-F4, M3d28-F1*, M3-d28-F2, M3d28-F3
   ⇒ Note: M3d28-F1* is identical to the fragment identified in M3-1 (Supplementary Text S2)
2. *De novo* alignment of unused reads from the Geneious alignment, searching specifically for unique contigs covering the duplication junction
   ⇒ Further evidence for the nine proposed duplication fragments above
3. Close observation of coverage data from the Geneious alignment (main manuscript Figure 8)
   ⇒ Regions of further interest identified
4. Manual inspection of regions of interest in Geneious alignment
   ⇒ Identification of 2 further predicted fragments: M1d28-F2, M2d28-F1
5. Demonstration of significant numbers of raw reads aligning to the unique duplication junction proposed for each population (and 0 for the same alignment performed with the raw reads of other populations) (see Tables 3.1 and 3.2 below).

#### 3.2 Duplication junction sequences

This section lists the 11 predicted duplication junction sequences in the day 28 populations of lines M1, M2, and M3. These are the sequences to which the raw reads from the relevant sequencing sample were aligned to obtain the read numbers listed in Tables 3.1 and 3.2 below.

```
>M1d28-F1-junct (30 bp)
CGGCGCGCTCCCCGagTTCGGTGTTCGCGT

>M1d28-F2-junct (30 bp)
TGTCAGCAACTCTTTCCCATCATGCACACG

>M1d28-F3-junct (30 bp)
AACGCTTTTGCAGCAtGCGCGCCGCCTGGC

>M1d28-F4-junct (55 bp)
TACAATCTAACTGTCaccccgCGATCCAAATGTGGGAGGGGGCTTGCCCCCGATA

>M2d28-F1-junct (54 bp)
ATCCCCGATGGTTAcTCGCCACGGGGGTGGTGCGCTACATTGTCGGTAATCCCG

>M2d28-F2-junct (30 bp)
CCCACGCCACGGCATTTCGACGCTCACAAA
```

>M2d28-F3-junct (30 bp)
CAATCTCCACGGCGCCCGCCGCGCAGATGC

>M2d28-F4-junct (128 bp)
GGGTAACGGCCGCTATGGGTTCCGCTTTTACAGCGGCTCACTTTTGAAGagcgcaaaagtaagcaaaacgctctt
gccccaaCACTCGGCACCTCGCTCACGCTCGGTGTGCCCGTAATCCGACAGGG

>M3d28-F1-junct* (88 bp) (identical to M3-1_junct)
GCCGTCCACCTATCgTGGGAGGGGGCTTGCCCCCGATAGCGGTGTGTCAGTCGATTCATCAGGTGACTGATTCAG
CGCCATCGGGGCG

>M3d28-F2-junct (30 bp)
ACCGCCGAACGCTTCTGCCGAACAGGCCCT

>M3d28-F3-junct (30 bp)
AACCCCGGTGCCTGCCCGATTACCGGCGCT

Duplication junctions key:
SIDE 1 of each junction
SIDE 2 of each junction
bases occurring on both side 1 and side 2 of the junction (*i.e.*, repeats)
*Note 1: each side (+ grey lettered shared region, where present) is a unique sequence (or, in the case of M1d28-F4-junct side 1, present twice) in the wildtype genome sequence.
**Note 2: no inversions; both sides of each junction occur on the same strand, in the same direction.
***Note 3: if the sides are highly similar, underline indicates bases that differ between side.

| **Fragment** | **Duplication fragment size** | | | **Computational evidence** | | |
|---|---|---|---|---|---|---|
| | bp | #duplicated genes[a] | #duplicated tRNA genes[a] | Evidence type | *breseq* freq. | Reads[b] |
| M1d28-F1 | 767,013 | 670 | 1 | Marginal prediction breseq, Geneious alignments | 3.10% | 9 |
| M1d28-F2 | 490,582 | 427 | 1 | Geneious alignments | n/a | 5 |
| M1d28-F3 | 444,617 | 403 | 1 | Mutation prediction breseq, Geneious alignments | 2.00% | 25 |
| M1d28-F4 | 4,273 | 7 | 1 | Mutation prediction breseq, Geneious alignments | 6.50% | 35 |
| M2d28-F1 | 490,215 | 427 | 1 | Geneious alignments | n/a | 10 |
| M2d28-F2 | 318,344 | 278 | 1 | Marginal prediction breseq, Geneious alignments | 4.10% | 7 |
| M2d28-F3 | 285,798 | 256 | 1 | Marginal prediction breseq, Geneious alignments | 2.30% | 108 |
| M2d28-F4 | 3,631 | 6 | 1 | Mutation prediction breseq, Geneious alignments | 32.00% | 3 |
| M3d28-F1* | 489,709 | 426 | 1 | Mutation prediction breseq, Geneious alignments | 15.60% | 28 |
| M3d28-F2 | 434,971 | 360 | 1 | Mutation prediction breseq, Geneious alignments | 32.70% | 77 |
| M3d28-F3 | 2,170 | 3 | 1 | Mutation prediction breseq, Geneious alignments | 20.00% | 80 |

**Table 3.1. Features of the large-scale duplication fragments.** [a]Lists of duplicated genes, including tRNA genes, can be found in Supplementary Table S3. [b]The number of raw reads found spanning the entire, unique duplication junction fragment in the population of identification (Geneious alignment). Duplication junction sequences are listed above, and read numbers corrected for duplication junction sequence length are listed in Table 3.2 below.

| Junction name | Junction seq. length (bp) | Number of raw reads covering entire junction sequence with no mismatches | | | | |
|---|---|---|---|---|---|---|
| | | **M1d28** 275.3 | **M2d28** 220.9 | **M3d28** 227.2 | **W1d28** 207.9 | **SBW25** 287.3 |
| ***Duplication fragments/junctions identified in day 21 isolates*** | | | | | | |
| M1-1_30bp_junct | 30 | 0 | 0 | 0 | 0 | 0 |
| M2-1_46bp_junct | 46 | 0 | 0 | 0 | 0 | 0 |
| M3-1_88bp_junct* | 88 | 0 | 0 | **28 (82)** | 0 | 0 |
| M4-1_714bp_junct | 714 | Junction seq. too long for unbroken coverage by 150 bp reads | | | | |
| M5-1_75bp_junct | 75 | 0 | 0 | 0 | 0 | 0 |
| ***Duplication fragments/junctions identified in day 28 populations*** | | | | | | |
| M1d28-F1-junct | 30 | **9** | 0 | 0 | 0 | 0 |
| M1d28-F2-junct | 30 | **5** | 0 | 0 | 0 | 0 |
| M1d28-F3-junct | 30 | **25** | 0 | 0 | 0 | 0 |
| M1d28-F4-junct | 55 | **35 (64)** | 0 | 0 | 0 | 0 |
| M2d28-F1-junct | 54 | 0 | **10 (18)** | 0 | 0 | 0 |
| M2d28-F2-junct | 30 | 0 | **7** | 0 | 0 | 0 |
| M2d28-F3-junct | 30 | 0 | **108** | 0 | 0 | 0 |
| M2d28-F4-junct | 128 | 0 | **3 (13)** | 0 | 0 | 0 |
| M3d28-F1-junct* | 88 | 0 | 0 | **28 (82)** | 0 | 0 |
| M3d28-F2-junct | 30 | 0 | 0 | **77** | 0 | 0 |
| M3D28-F3-junct | 30 | 0 | 0 | **80** | 0 | 0 |

**Table 3.2. Coverage of the predicted duplication junctions across various samples.** Each predicted duplication junction sequence (first column) was used as a reference to which the raw reads obtained during genome re-sequencing of nine populations were aligned (mean genomic coverage of each sample is provided under the population name). Numbers of raw reads aligning perfectly (no mismatches and unbroken coverage) to each duplication junction sequence, for each population, is listed. Numbers in parentheses are read numbers corrected for length of duplication fragment, relative to the usual 30 bp, to nearest whole read number). **Blue bold**=non-zero read numbers (notably, these occur exclusively in the population in which the duplication junction was identified; matches are entirely absent in the other populations). *these two junctions are identical. This result provides evidence that each of the predicted duplication junctions (and therefore, duplication fragments) is distinct from those predicted in other, independent populations.

The above results support the co-existence of 11 distinct, variously sized duplication fragments in the three mutant day 28 populations (4 fragments in M1-d28, 4 fragments in M2-d28, 3 fragments in M3-d28). Notably, there may still be other, undetected fragments – particularly those occurring between long (>150-bp) perfect repeats – present in some mutant day 28 populations.

#### Supplementary Text S4
#### Reference list of 42 unique tRNA sequences in *P. fluorescens* SBW25

GtRNAdb 2.0 (Chan and Lowe 2009; Chan and Lowe 2016) identifies 67 tRNA genes in *P. fluorescens* SBW25. These include 42 unique primary tRNA gene sequences (listed below). One (11_Cys-GCA-2-1) is a partial tRNA gene for which no contribution to the mature tRNA pool (i) is expected, or (ii) has been detected in our previous laboratory experiments (Ayan et al. 2020). However, as a control, the Cys-GCA-2-1 sequence was included in the alignment process.

These 42 sequences are used as references to align the YAMAT-seq data in this work. The raw YAMAT-seq data is available at NCBI GEO (GSE196705); a summary of the alignment results is provided in Supplementary Table S4, and an overview of pairwise tRNA expression analyses by DESeq2 are provided in Supplementary Table S5.

```
>1_Ala-GGC-1-2 76 bp Sc: 75.2
GGGGCTATAGCTCAGCTGGGAGAGCGCTTGCATGGCATGCAAGAGGtCAACGGTTCGATCCCGTTTAGCTCCACCA
>2_Ala-TGC-1-5 76 bp Sc: 82.7
GGGGCCATAGCTCAGCTGGGAGAGCGCCTGCCTTGCACGCAGGAGGtCAACGGTTCGATCCCGTTTGGCTCCACCA
>3_Arg-ACG-1-2 77 bp Sc: 84.0
GCACTCGTAGCTCAGCTGGAtAGAGTACTCGGCTACGAACCGAGCGGtCACAGGTTCGAATCCTGTCGAGTGCACCA
>4_Arg-CCG-1-1 77 bp Sc: 83.6
GCATCCGTAGCTCAGCTGGAtAGAGTACTGCCCTCCGAAGGCAGGGGtCGTGGGTTCGAATCCCGCCGGGTGCACCA
>5_Arg-CCT-1-1 77 bp Sc: 72.4
GTCCCAGTAGCTCAATTGGAtAGAGCATCCCCCTCCTAAGGGGAAGGtTGGCCGTTCGAACCGGCCCTGGGACACCA
>6_Arg-TCT-1-1 77 bp Sc: 89.3
GCGCCCGTAGCTCAGCTGGAtAGAGCATCCGCCTTCTAAGCGGATGGtCGCAGGTTCGAGTCCTGCCGGGTGCGCCA
>7_Asn-GTT-1-1 76 bp Sc: 80.6
TCCGTGATAGCTCAGTCGGTAGAGCAAATGACTGTTAATCATTGGGtCCCAGGTTCGAGTCCTGGTCACGGAGCCA
>8_Asn-GTT-2-1 76 bp Sc: 77.6
TCCGCGATAGCTCAGTTGGTAGAGCAAATGACTGTTAATCATTGGGtCCCTGGTTCGAGTCCAGGTCGTGGAGCCA
>9_Asp-GTC-1-4 77 bp Sc: 90.7
GCAGCGGTAGTTCAGTCGGTtAGAATACCGGCCTGTCACGCCGGGGGtCGCGGGTTCGAGTCCCGTCCGCTGCGCCA
>10_Cys-GCA-1-1 74 bp Sc: 65.3
GGCCGAGTAGCAAAATGGTTATGCAGCGGATTGCAAATCCGCCTaCGCCGGTTCGATTCCGACCTCGGCCTCCA
>11_Cys-GCA-2-1 112 bp Sc: 23.4 (tRNA fragment)
GAGTAAATGTTGGTgAtgcggaatagattatcatataacttattgaaaataatgaggaTATTCGTGGATTGCAAATCCGCCTa
CGCCGGTTCGATTCCGACCTCGGCCTCCA
>12_Gln-TTG-1-1 75 bp Sc: 69.7
AGGGGCGTCGCCAAGCGGTAAGGCAGCAGGTTTTGATCCTGCCATgCGTTGGTTCGAATCCAGCCGCCCCTGCCA
>13_Glu-TTC-1-4 76 bp Sc: 68.9
GTCCCCTTCGTCTAGTGGCctAGGACACCGCCCTTTCACGGCGGTAaCAGGGGTTCGAGTCCCCTAGGGGACGCCA
>14_Gly-CCC-1-1 74 bp Sc: 74.9
GCGGGTATAGTTTAATGGTAGAACAGTAGCTTCCCAAGCTTCCGaCGAGGGTTCGATTCCCTCTACCCGCTCCA
>15_Gly-GCC-1-3 76 bp Sc: 88.4
GCGGGAATAGCTCAGTTGGTAGAGCACGACCTTGCCAAGGTCGGGGtCGCGAGTTCGAGTCTCGTTTCCCGCTCCA
>16_Gly-TCC-1-1 74 bp Sc: 83.2
GCGGGTATAGTTTAGTGGTAGAACCTCAGCCTTCCAAGCTGATGaTGCGGGTTCGATTCCCGCTACCCGCTCCA
>17_His-GTG-1-2 76 bp Sc: 73.3
GTGGGCGTAGCTCAGTTGGTAGAGCACGGGATTGTGACTCCCGTTGtCGAGGGTTCGATCCCCTTCGTCCACCCCA
>18_Ile-GAT-1-5 77 bp Sc: 88.9
GGGTCTGTAGCTCAGTTGGTtAGAGCGCACCCCTGATAAGGGTGAGGtCGGCAGTTCGAATCTGCCCAGACCCACCA
>19_Ile2-CAT-1-1 77 bp Sc: 89.4
GGGCCTATAGCTCAGTTGGTtAGAGCAGGGGACTCATAATCCCTTGGtCGTAGGTTCGAGTCCTACTGGGCCCACCA
>20_Leu-CAA-1-1 85 bp Sc: 70.8
GCCCTGATGGCGGAATTGGTaGACGCGGCGGATTCAAAATCCGTTTTCGAAAgGAGTGGGAGTTCGAGTCTCCCTCGGGGCAC
CA
>21_Leu-CAG-1-2 87 bp Sc: 71.6
GCCCAGGTGGTGAAATTGGTaGACACGCCAGCTTCAGGTGCTGGTGATCGCAAGGTCGTGGAAGTTCGAGTCTTCTCCTGGGC
ACCA
```

```
>22_Leu-GAG-1-1 86 bp Sc: 60.3
GCCGAGGTGGTGGAATTGGTaGACACGCAACCTTGAGGTGGTTGTGCCCATAGGGTgTAGGGGTTCGAGTCCCCTTCTCGGTA
CCA
>23_Leu-TAA-1-1 87 bp Sc: 71.0
GCCCGAATGGCGAAACTGGTaGACGCATGGGACTTAAAATCCCCCGCTCGTAAGGGCGTCCCGGTTCGATTCCGGGTTCGGGC
ACCA
>24_Leu-TAG-1-1 85 bp Sc: 70.1
GCGGATGTGGTGGAATTGGTaGACACACTGGATTTAGGTTCCAGCGCCGCGAGGCGTAAGAGTTCGAGTCTCTTCATCCGCAC
CA
>25_Lys-TTT-1-2 76 bp Sc: 86.5
GGGTCGTTAGCTCAGTTGGTAGAGCAGTTGGCTTTTAACCAATTGGtCGTAGGTTCGAATCCCACACGACCCACCA
>26_Met-CAT-1-1 77 bp Sc: 77.5
GGCTACATAGCTCAGTTGGTtAGAGCATAGCATTCATAATGCTGGGGtCCGGGGTTCAAGTCCCTGTGTAGCCACCA
>27_Phe-GAA-1-1 76 bp Sc: 81.1
GCCCAGATAGCTCAGTCGGTAGAGCAGGGGATTGAAAATCCCCGTGtCGGCGGTTCGATTCCGTCTCTGGGCACCA
>28_Pro-GGG-1-1 77 bp Sc: 67.6
CGGGGCGTAGCGCAGTCcGGTAGCGCACTAGCATGGGGTGCTAGGGGtCGAGTGTTCGAATCACTCCGTCCCGACCA
>29_Pro-TGG-1-2 77 bp Sc: 73.4
CGGGGTATAGCGCAGTCcGGTAGCGCGCCTGCTTTGGGAGCAGGATGtCAGGAGTTCGAATCCCCTTACCCCGACCA
>30_Ser-CGA-1-1 90 bp Sc: 71.0
GGAGAGATGCCAGAGTGGCcgAATGGGACGGATTCGAAATCCGTTGTACCTTCACCGGTACCTAGGGTTCGAATCCCTATCTC
TCCGCCA
>31_Ser-GCT-1-1 91 bp Sc: 75.0
GGAGAGCTGGCCGAGTGGCcgAAGGCGCTCCCCTGCTAAGGGAGTACACCTCAAaAGGGTGTCGGGGGTTCGAATCCCCCGTT
CTCCGCCA
>32_Ser-GGA-1-1 90 bp Sc: 75.7
GGTGAAGTGTCCGAGTGGCttAAGGAGCACGCCTGGAAAGTGTGTATACAAGAAATTGTATCGAGAGTTCGAATCTCTCCTTC
ACCGCCA
>33_Ser-TGA-1-1 91 bp Sc: 75.4
GGGAAATTGGCAGAGTGGTtgAATGCACCGGTCTTGAAAACCGGCGGACGTTAAtAGCGTCTCCAGGGTTCGAATCCCTGGTT
TCCCGCCA
>34_Thr-CGT-1-1 73 bp Sc: 69.7
GCCCGTGTAGCTCAGTCGGTAGAGCAGCGCACTCGTAACGCGAAGGtCGCAGGTTCGATTCCTGTCTCGGGCACCA
>35_Thr-GGT-1-1 76 bp Sc: 80.7
GCTCTTGTAGCTCAGTTGGTAGAGCACACCCTTGGTAAGGGTGAGGtCAGCGGTTCAAATCCGCTCAAGAGCTCCA
>36_Thr-TGT-1-1 76 bp Sc: 87.9
GCCGGTATAGCTCAGTTGGTAGAGCAACTGACTTGTAATCAGTAGGtCCCGGGTTCGACTCCTGGTGCCGGCACCA
>37_Trp-CCA-1-1 76 bp Sc: 86.5
AGGTCAGTAGCTCAATTGGCAGAGCGACGGTCTCCAAAACCGTAGGtTGGGGGTTCGATTCCCTCCTGACCTGCCA
>38_Tyr-GTA-1-1 85 bp Sc: 77.1
GGAGGGGTTCCCGAGCGGCcaAAGGGATCAGACTGTAAATCTGACGTCTACGACTtCGAAGGTTCGAATCCTTCCCCCTCCAC
CA
>39_Val-GAC-1-1 77 bp Sc: 78.1
AGGCACGTAGCTCAGTTGGTtAGAGCACCACCTTGACATGGTGGGGGtCGTTGGTTCGAGTCCAATCGCGCCTACCA
>40_Val-TAC-1-3 76 bp Sc: 84.4
GGGTGATTAGCTCAGCTGGGAGAGCATCTGCCTTACAAGCAGAGGGtCGGCGGTTCGATCCCGTCATCACCCACCA
>41_fMet-CAT-1-1 77 bp Sc: 78.4
CGCGGGGTGGAGCAGTCtGGTAGCTCGTCGGGCTCATAACCCGAAGGtCGTCGGTTCAAATCCGGCCCCCGCAACCA
>42_fMet-CAT-2-2 77 bp Sc: 76.7
CGCGGGATGGAGCAGTCtGGTAGCTCGTCGGGCTCATAACCCGAAGGtCGTCGGTTCAAATCCGGCTCCCGCAACCA
```

### Supplementary Figures

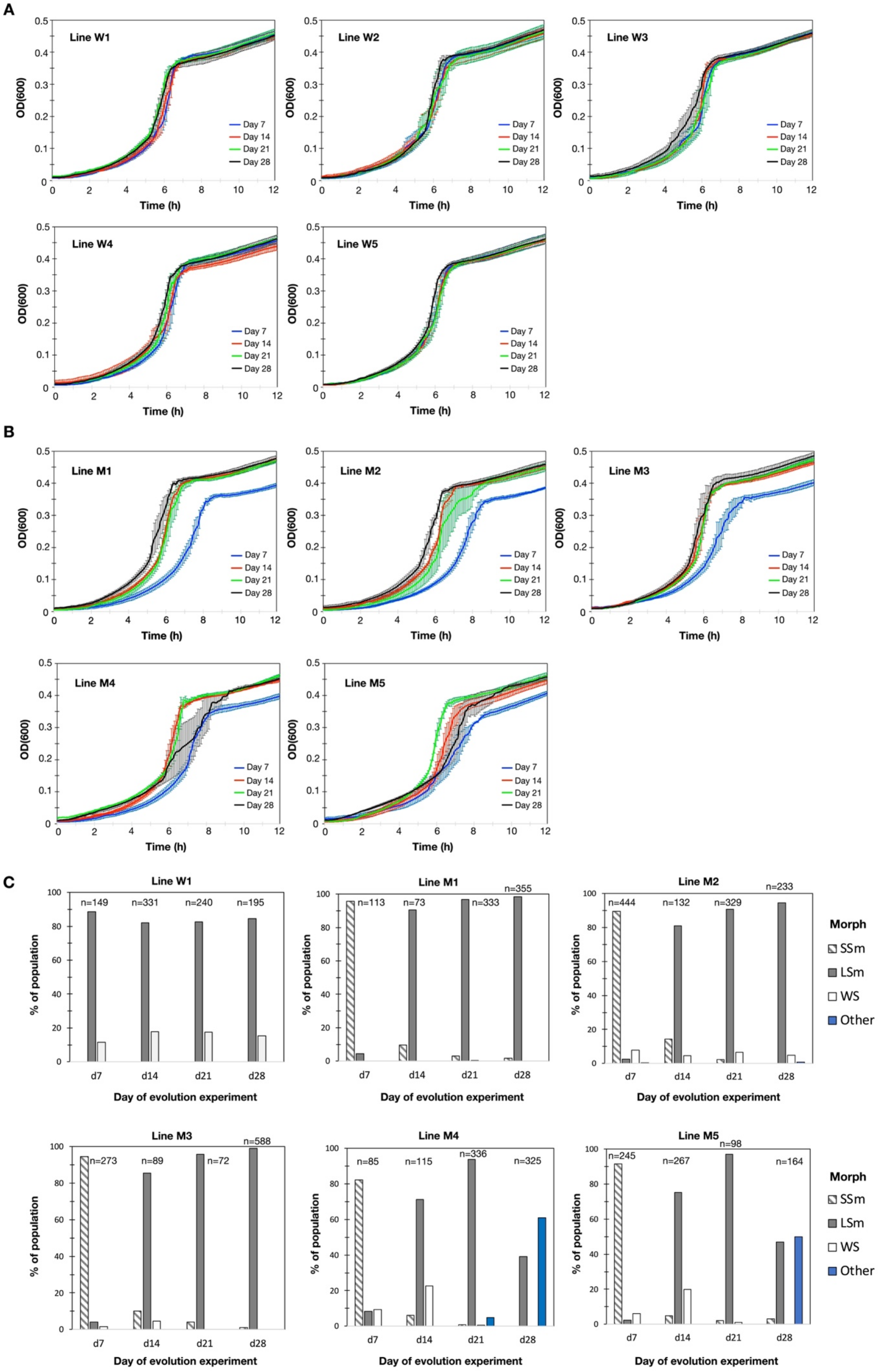

**Supplementary Figure S1. Phenotypic analyses of evolving populations on days 7, 14, 21, 28.** Growth curves in KB medium for all wildtype (**A**) and ΔEGEG (**B**) evolutionary lines. Wildtype populations show little change in growth profiles; line W1 was used as a representative control from this point forward. Mutant populations show general growth improvement, with the first increase typically occurring between days 7 and 14. A second increase can be observed between days 21 and 28 of lines M1 and M2 (from which ~1 Mb duplication fragments were isolated on day 21; see Table 1). Lines M4 and M5 show an unexpected decrease in growth on day 28 (see panel C). Per graph, each line represents the mean ± 1 SE of 10 (line M3 day 28) or 12 replicates (see main Methods). (**C**) SBW25 is known to give rise to morphologically distinct colony morphotypes, many of which are underpinned by well-characterised mutations (*e.g.,* Rainey and Travisano 1998; Gallie et al. 2015). A morphological analysis of colonies from populations W1, M1-M5 reveals rapidly changing colony morphologies. SSm=small smooth, LSm=large smooth, WS=wrinkly spreader, "Other"=any other phenotype. Consistent with the non-canonical growth profiles in panel B, many "other" colonies were observed on days 21-28 of lines M4 and M5. 16S rRNA Sanger sequencing shows these to be non-SBW25 contaminants. Lines M4 and M5 were excluded from downstream population analyses.

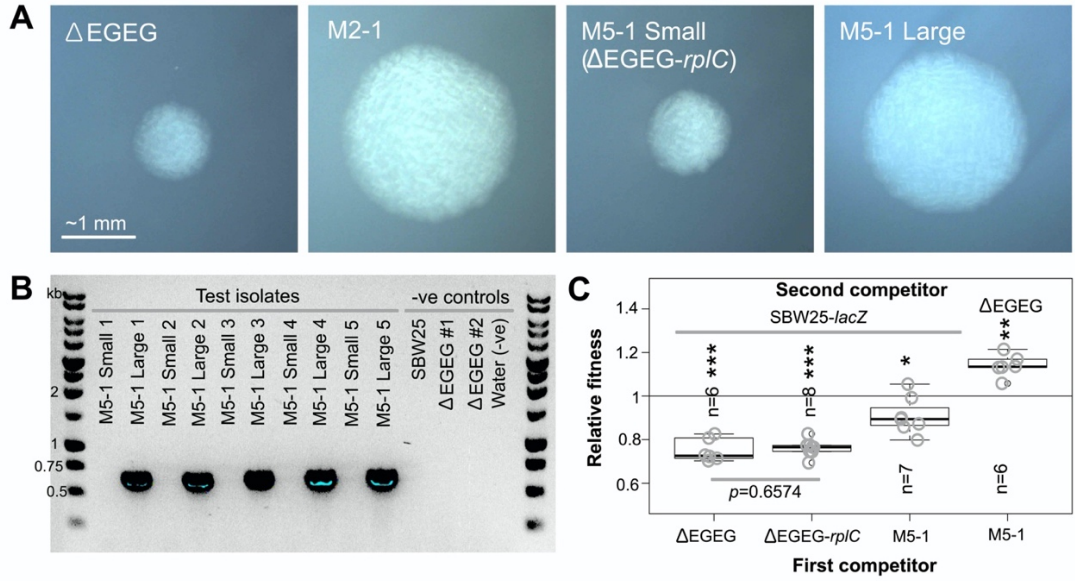


**Supplementary Figure S2. A SNP in *rplC* (c128t, causing amino acid change T43I) does not contribute significantly to the increased fitness of M5-1.** (**A**) When grown in overnight KB culture, M5-1 (carrying the ~1 Mb duplication fragment M5f, and a SNP in *rplC*) produces a mixture of large and small colonies (see also Figure 6). These two colony types are photographed here, along with ΔEGEG (carrying neither mutation) and M2-1 (carrying only the ~1 Mb duplication fragment). Colonies were grown on KB agar at 28˚C for 30 hours, and photographed under the same magnification (scale bar applies to all images). The exposure of some images was uniformly altered to increase visibility. (**B**) Five single (large) M5-1 colonies were each inoculated into 4 ml liquid KB, incubated at 28˚C overnight (shaking), and plated on KB agar. After 30 hours' incubation (28˚C), one small and one large colony was purified from each of the five sets of plates. A duplication junction PCR demonstrated that 611 bp surrounding the emergent M5-1 duplication junction can be amplified from all five large colonies, but none of the small colonies (and none of the controls). PCR and Sanger sequencing of *rplC* demonstrated that all ten isolates retained the *rplC* mutation. M5-1 small colony #1 – which carries only the *rplC* mutation, and no duplication fragment – was named ΔEGEG-*rplC* and used in the subsequent competition experiment. (**C**) Direct, 1:1 competition experiments were performed in liquid KB for 24 hours (28˚C, shaking). Between six and eight replicate competitions were performed for each strain pair. Box plots of the relative fitness of competitor 1 (*x*-axis) and competitor 2 (top). Relative fitness >1 means competitor 1 wins, <1 means competitor 2 wins. Both ΔEGEG and ΔEGEG-*rplC* were outcompeted by SBW25-*lacZ* (parametric one-sample *t*-tests $p$<8.2e-05), and no difference in relative fitness values was detected between the two competitions (parametric two-sample *t*-test $p$=0.6574). Datapoints from all replicates are overlaid on boxplots (grey circles), and ***$p$<0.001, **$p$<0.01, *p<0.05. Data from competitions 1, 3 and 4 are also reported in Figure 3.

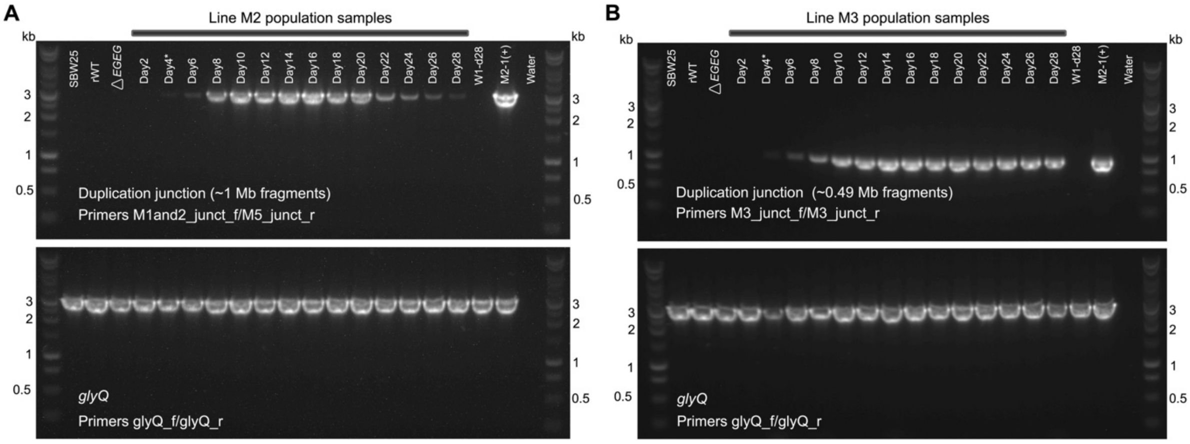


**Supplementary Figure S3. Examples of the agarose gels used to demonstrate rapid evolutionary dynamics of specific duplication fragments in mutant lines M2 and M3.** (**A**) The dynamics of the set of ~1 Mb duplication fragments captured by primer pair M1and2_junct_f/M5_junct_r, across evolving line M2. (**B**) The dynamics of the set of ~0.49 Mb duplication fragments captured by primer pair M3_junct_f/r, across evolving line M3. For both panel A and panel B, duplication junction PCRs were performed as outlined in the Methods. Briefly: genomic DNA was isolated from each population (or, in the case of controls, genotype), and 20 ng of each gDNA preparation was used for each of two PCRs: (PCR-1; top gels) targeting the duplication junction of interest (top gels), and (PCR-2; bottom gels) targeting a housekeeping region outside the genomic region where the duplication events occur (*glyQ*, at ~0.01 Mb of SBW25 genome). The presence and/or intensity of the duplication junction PCR product is expected to change across the evolving population (as duplication fragments arise and spread), while that of the housekeeping region is expected to remain constant. The intensity of each duplication junction PCR product was measured in imageJ2, and normalized against that of the corresponding housekeeping PCR (PCR-2; see Supplementary Table S6). The entire process – including gDNA preparation – was repeated three times for each set of PCRs. The gels shown are for PCR round 3 in line M2 (panel A) and PCR round 2 in line M3 (panel B). The complete set of gels and analyses can be found in Supplementary Table S6, and final results are presented in Figure 7A-B.

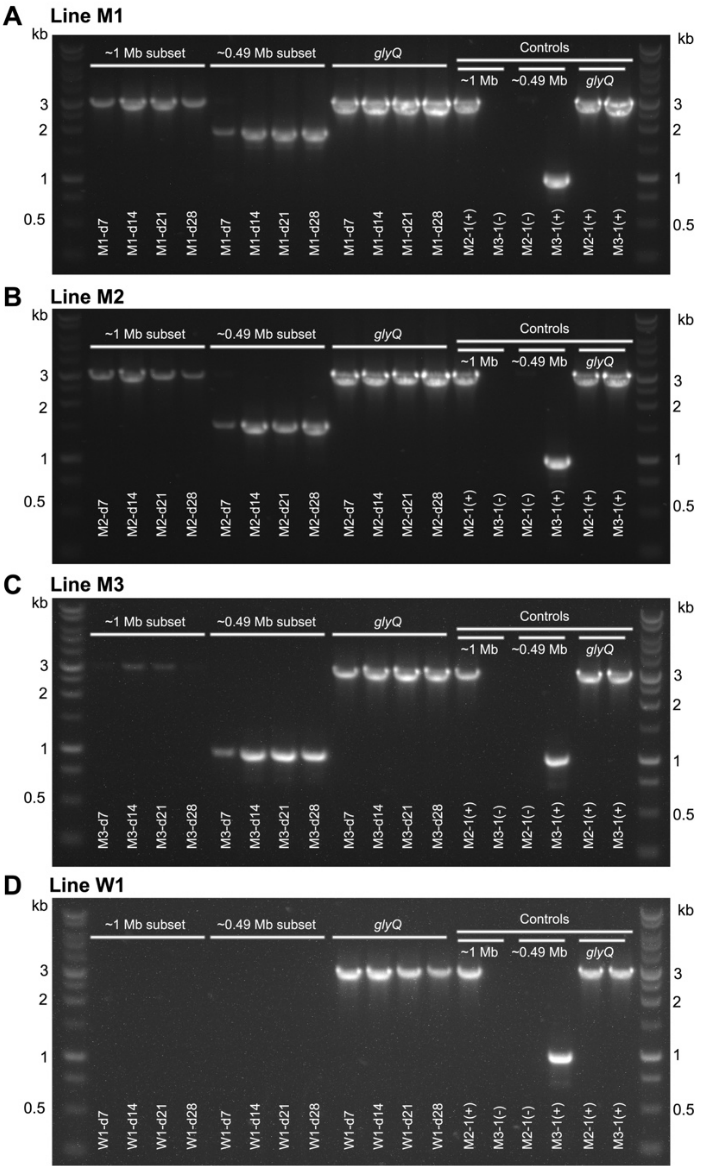


**Supplementary Figure S4. Examples of the agarose gels used to demonstrate the presence of multiple duplication fragments per evolving mutant line.** The dynamics of ~1 Mb and ~0.49 Mb duplication fragment sets across evolving mutant lines M1 (**A**), M2 (**B**), M3 (**C**) and a representative control line (W1, **D**). Duplication junction PCRs were performed as outlined in the Methods. Briefly: gDNA was isolated from samples of interest. 20 ng of each gDNA preparation was used for each of three PCRs (left to right per gel): (i) PCR-1 targets the ~1 Mb subset of duplication fragments captured by primer pair M1and2_junct_f/M5_junct_r, (ii) PCR-2 targets the ~0.49 Mb subset captured by primer pair M3_junct_f/r, and (iii) PCR-3 targets the housekeeping region *glyQ* (outside of the region affected by duplication events). PCR products representing both the ~1 Mb and ~0.49 Mb duplication fragment subsets are detected in all three mutant lines (and not the control line W1). Notably, the size of the PCR product indicating the ~0.49 Mb fragment set differs by line (line M1 ~1.8 kb; line M2 ~1.5 kb; line M3 ~1 kb), indicating duplication fragments with subtly different endpoints in each line (see also Supplementary Figure S5B). The intensity of each duplication junction PCR product was measured in imageJ2, and normalized against that of the corresponding housekeeping PCR (PCR-3; see Supplementary Table S6). The entire process – including gDNA preparation – was repeated three times for each set of PCRs; only one example gel is shown for each evolving line. The complete set of gels and analyses can be found in Supplementary Table S6 and Supplementary Figure S5A. Final results are presented in Figure 7C.

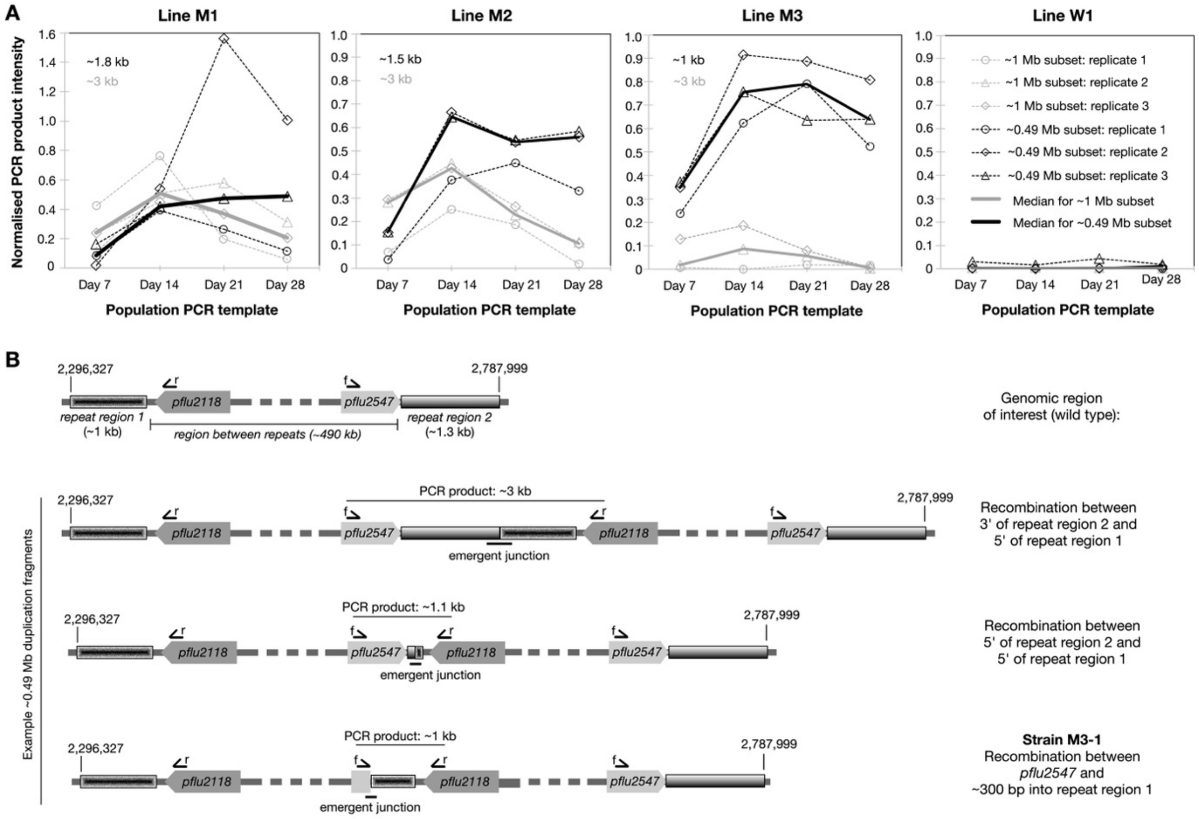


**Supplementary Figure S5. Each evolving mutant line contains multiple types of competing duplication fragments.** (**A**) Three independent replicates of the normalized intensity of duplication junction PCR products from the ~1 Mb duplication fragment set (dotted grey lines) and the ~0.49 Mb duplication fragment set (dotted black lines) across evolving lines M1, M2, M3, W1. Solid lines indicate the respective medians (which are also presented in Figure 7C). The approximate size of the respective PCR product is indicated in grey and black in the top left corner of each gel. Notably, the product of the PCR targeting the ~0.49 Mb duplication fragment set (in black) is unique to each evolving line, indicating subtly different duplication fragments. Sanger sequencing of the ~1.8 kb, ~1.5 kb, and ~1 kb PCR products confirms that the 5' and 3' ends of these PCR products show homology to positions ~2.3 Mb and ~2.8 Mb of the SBW25 chromosome (*i.e.*, result from the tandem duplication between these positions). (**B**) Cartoon depiction of the mechanism behind the generation of ~0.49 Mb duplication fragments with distinct PCR product sizes. The ~0.49 Mb duplication fragments typically arise by unequal recombination between two repetitive regions of the SBW25 genome: a ~1 kb intergenic region at position ~2.3 Mb and a ~1.8 kb intergenic region at position ~2.8 Mb (compensatory *glyGCC* lies between these repeats, at position ~2.4 Mb) (top). Unequal recombination can, hypothetically, occur between these regions at any number of different points, resulting in PCR products of between ~3 kb and ~1.1 kb (with primer pair M3_junct_f/r; see top two example duplication fragments). The slightly smaller 1 kb PCR product in line M3 has been demonstrated to result from a duplication of the region between *pflu2547* and the intergenic region at position ~2.8 Mb (confirmed by Sanger sequencing; and also isolate M3-1 in Table 1). This means that, while homology between endpoints is likely to increase the rate of recombination, it is not a requirement for tandem duplication events.

**Supplementary Tables (legends)**

This section contains the legends for all supplementary tables available as excel files.

**Supplementary Table S1**. Details of genotypes, plasmids, oligonucleotides, and duplication junctions in this study.

**Supplementary Table S2**. **List of mutations predicted from the whole genome re-sequencing of isolates from day 21 of the evolution experiment.** Illumina whole genome sequence data were obtained for six isolates from day 21 of the evolution experiment: W1-1 (derived from representative wildtype control line W1), M1-1, M2-1, M3-1, M4-1, and M5-1 (each derived from an independent *EGEG* deletion line). This file provides a summary and a full report of the mutations predicted in each isolate. In addition to the large tandem duplications detailed in Table 1, one unique point mutation was identified: M5-1 carries a non-synonymous point mutation in *rplC* (encoding 50S ribosomal protein L3). Some putative mutations were identified in multiple isolates, many of which have also been identified in the ancestral SBW25 wildtype that was sequenced as part of the population sequencing later in this manuscript (see Supplementary Table S7). Hence, these predicted mutations are likely to either be present in the starting strain, or constitute computational alignment errors. As such, these putative mutations are not expected to be relevant for the described fitness effects.

**Supplementary Table S3**. **List of genes contained in each duplication fragment.** The spreadsheet lists SBW25 gene annotations from NCBI (6,176 genes; left of the spreadsheet), followed by lists of genes that are duplicated in the 15 identified duplication fragments (left to right). For ease of comparison, the duplication details for each fragment are provided on the same line numbers as the SBW25 gene list (*i.e.*, for each fragment, scroll down to the first duplicated gene). The smallest number of complete genes in a single fragment is 3: fragment M3d28-F3 contains *pgsA* (phospholipid biosynthesis), *glyGCC* (tRNA), *pflu2192* (predicted pseudogene). Only two genes, *glyGCC* and *pflu2192*, are duplicated in every fragment; these 'core set' genes are highlighted in cyan.

**Supplementary Table S4. YAMAT-seq data showing changes in mature tRNA pools due to engineering and/or evolution.** The first tab contains index details and a summary of the raw YAMAT-seq reads (GEO accession number GSE196705) (Edgar et al. 2002); for each of 24 samples (three replicates of eight genotypes): a minimum of ~1.46 million reads of the expected size (80-151 bp) was obtained per sample. In each case, between 96.7 % and 98.0 % of these aligned to the list of 42 reference SBW25 tRNA sequences (provided in Supplementary Text S4). The subsequent eight tabs contain the YAMAT-seq data for each of the nine genotypes tested (SBW25, rWT, ΔEGEG, Δ*glyGCC*, W1-1, M1-1, M3-1, M5-1). Each tab contains (i) the numbers of reads for 42 reference tRNAs, for three replicates (left columns), (ii) numbers of reference reads for 39 tRNA species in

SBW25 (*e.g.*, tRNA-Asn-GTT is the sum of reference sequences 7_Asn-GTT-1-1 and 8_Asn-GTT-2-1; middle columns), (iii) the proportion of each tRNA species in the mature tRNA pool for each of the three samples (right columns), and (iv) a scatter plot of the YAMAT-seq proportions versus the proportion of the tRNA gene set encoding the tRNA species. Magenta=tRNA-Gly(GCC), blue=tRNA-Glu(UUC).

**Supplementary Table S5. Statistical analysis to detect differences in the expression of 39 tRNA species between pairs of strains.** This file contains the DESeq2 values from the YAMAT-seq analysis used in Figure 5. Using the aligned YAMAT-seq read data (Supplementary Table S4) as input, DESeq2 output consists of: baseMean1 and baseMean2 (the normalized mean expression level of three replicates of strain 1 and strain 2); fold.change1 and fold.change2 (fold change calculated by baseMean1/baseMean2 or baseMean2/baseMean1, respectively); log2.fold.change1 and log2.fold.change2 (log2 of fold.change1 or log2 of fold.change2, respectively); *p*value (calculated by assuming a binomially distributed read coverage analogous to Fisher's exact test (Robinson and Smyth 2008; Anders and Huber 2010; Anders et al. 2015)); padj (*p*value adjusted for multiple testing with the Benjamini-Hochberg procedure, which controls for false discovery rate (Anders and Huber 2010; Anders et al. 2015)). For each pairwise comparison, rows (tRNA species) are ordered according to increasing padj. All rows with a padj<0.01 (above the solid black line) are then ordered by decreasing log2(fold.change1), and those with a negative log2(fold.change1) value are ordered according to decreasing fold.change2. This places tRNA species with statistically significant differences in expression first, with those higher in genotype 1 listed at the top (ordered by decreasing size of expression difference; pink), followed by those higher in genotype 2 (ordered by decreasing expression difference; green). The first tab contains comparisons investigating the effect of deleting the tRNA gene quad (*gluTTC-glyGCC-gluTTC-glyGCC*; *EGEG*) and the lone *glyGCC* gene copy. The second tab contains comparisons investigating the effect of mutations acquired during the evolution experiment (duplication fragments and/or SNPs). The third tab contains the comparisons between each of the five duplication genotypes and SBW25. The fourth tab contains comparisons between different evolved isolates.

**Supplementary Table S6. The dynamics of large-scale duplication fragments in evolving populations.** This file contains the gel images, intensity measurements, and normalization calculations from the duplication junction PCRs in Figure 7 and Supplementary Figure S5.

**Supplementary Table S7. Details of the whole genome re-sequencing of populations from day 28 of the evolution experiment.** Illumina whole genome sequencing was obtained for nine populations: day 28 M1-M3 and W1-W5, and an ancestral SBW25 genotype (treated as a population sample as a control for the downstream computational analyses). This file contains the details of the raw sequencing reads and the *breseq* polymorphism output, for each sample.
